# Multifunctional ferromagnetic fiber robots for navigation, sensing, and treatment in minimally invasive surgery

**DOI:** 10.1101/2023.01.27.525973

**Authors:** Yujing Zhang, Xiaobo Wu, Ram Anand Vadlamani, Youngmin Lim, Jongwoon Kim, Kailee David, Earl Gilbert, You Li, Ruixuan Wang, Shan Jiang, Anbo Wang, Harald Sontheimer, Daniel English, Satoru Emori, Rafael V. Davalos, Steven Poelzing, Xiaoting Jia

## Abstract

Small-scale robots capable of remote active steering and navigation offer great potential for biomedical applications. However, the current design and manufacturing procedure impede their miniaturization and integration of various diagnostic and therapeutic functionalities. Here, we present a robotic fiber platform for integrating navigation, sensing, and therapeutic functions at a submillimeter scale. These fiber robots consist of ferromagnetic, electrical, optical, and microfluidic components, fabricated with a thermal drawing process. Under magnetic actuation, they can navigate through complex and constrained environments, such as artificial vessels and brain phantoms. Moreover, we utilize Langendorff mouse hearts model, glioblastoma microplatforms, and in vivo mouse models to demonstrate the capabilities of sensing electrophysiology signals and performing localized treatment. Additionally, we demonstrate that the fiber robots can serve as endoscopes with embedded waveguides. These fiber robots provide a versatile platform for targeted multimodal detection and treatment at hard-to-reach locations in a minimally invasive and remotely controllable manner.

## Introduction

Small-scale robotic devices capable of remotely navigating through complex and dynamic environments are arising as promising technology for biomedical applications^1–4^. Owing to their flexibility and steerability, these robotic devices can potentially offer minimally invasive, localized, and targeted diagnostic and therapeutic procedures, for next-generation percutaneous coronary intervention (PCI), atrial fibrillation (AF) ablation, gastrointestinal endoscopy, brain surgery, etc.^5–10^, where the operating space is confined. Despite the great potential of biomedical robotic devices, several challenges and critical issues exist which hinder their translation to clinics, including (1) difficulty in scaling robotic devices down to the micrometer scale, limiting the types of lesions that can be accessed^4,11,12^, (2) inefficiency and inaccuracy of the guidance and navigation process, impeding the delivery of localized and precise therapy deep inside the body^13^, and (3) the lack of integrated multimodal sensing and treatment systems, restricting the functions that can be achieved via robotic devices^3,4,6^.

To address the first two challenges, researchers have recently developed ferromagnetic soft robots composed of flexible polymer matrices with doped ferromagnetic micro/nanoparticles^14–18^. The response of the ferromagnetic soft robots to external magnetic fields can be precisely predicted and designed by calculating the generated torques or forces using quantitative models^19,20^. As the actuation relies on the dispersed ferromagnetic micro/nanoparticles, these robotic devices can be miniaturized and encoded on a microscale^21–24^, which makes them a promising approach for minimally invasive surgery.

Despite the advantages offered by the ferromagnetic soft robots, a major challenge in these devices is the lack of multimodal diagnostic and therapeutic functions. Recently, several attempts have been made to enable multimodal capabilities in microscale robotics. Conventional clean-room technology allows for the fabrication of micro-robotic probes with integrated electronic components for heating and flow sensing^25^. However, the total length of these devices is limited by the size of the silicon wafer, well below the length requirement for most interventional surgeries. In addition, the fabrication involves complicated and costly procedures. Injection molding can be used to incorporate simple components such as an optical fiber or a hollow channel into robots^21,26^, but is still hard to realize complex multimaterial structures with a micro-scale resolution. So far, scalable submillimeter ferromagnetic robots with multiple diagnostic and therapeutic functions have yet to be developed.

Over the past few years, the thermal drawing process (TDP) has been rapidly evolving as a powerful tool to fabricate scalable multimaterial fibers for biomedical applications. The TDP allows the fabrication of hundreds of meter-long and highly uniform multimaterial fibers with complex architectures and textures to achieve multiple functionalities^27^. Examples include multimaterial fibers for electrophysiological and chemical sensing^28–30^, nerve and skeletal muscle regeneration^31,32^, and optical and chemical modulation^33–35^. The ease of fabrication and capability of incorporating complex materials and structures inside fibers makes TDP an ideal candidate for manufacturing scalable ferromagnetic fiber robots with integrated diagnostic and therapeutic functions.

Here, we present submillimeter multifunctional ferromagnetic fiber robots (MFFRs) which integrate navigation, sensing, and treatment capabilities. Using TDP, we integrate ferromagnetic, electrical, optical, and microfluidic components in a single fiber which can be as long as hundreds of meters. With controlled external magnetic fields, these fiber robots can be steered through complex and constrained environments, such as tortuous vascular and brain phantoms. By integrating a functional core consisting of electrical, optical, and microfluidic components, we demonstrate the sensing and therapeutic capabilities of the fiber robots. In particular, we utilize a Langendorff-perfused mouse heart model to show that the fiber robots can enable intracardiac electrogram recording, pacing, and bioimpedance monitoring. The fiber robots can also deliver pulsed electric fields to irreversibly electroporate glioblastoma tumor cells in hydrogel platforms. Furthermore, we show that our fiber robots can record brain activities with a single unit resolution and optically modulate neural activities using *in vivo* mouse models. Additionally, we demonstrate that the fiber can be used for imaging with embedded waveguides. Given their miniaturized size and multifunctionality, our ferromagnetic fiber robots can not only improve existing surgeries, but also get access to and provide diagnosis and treatment in previously inaccessible lesions, as illustrated in **Fig.1A**, thereby significantly improving the surgical outcomes.

**Fig. 1.**
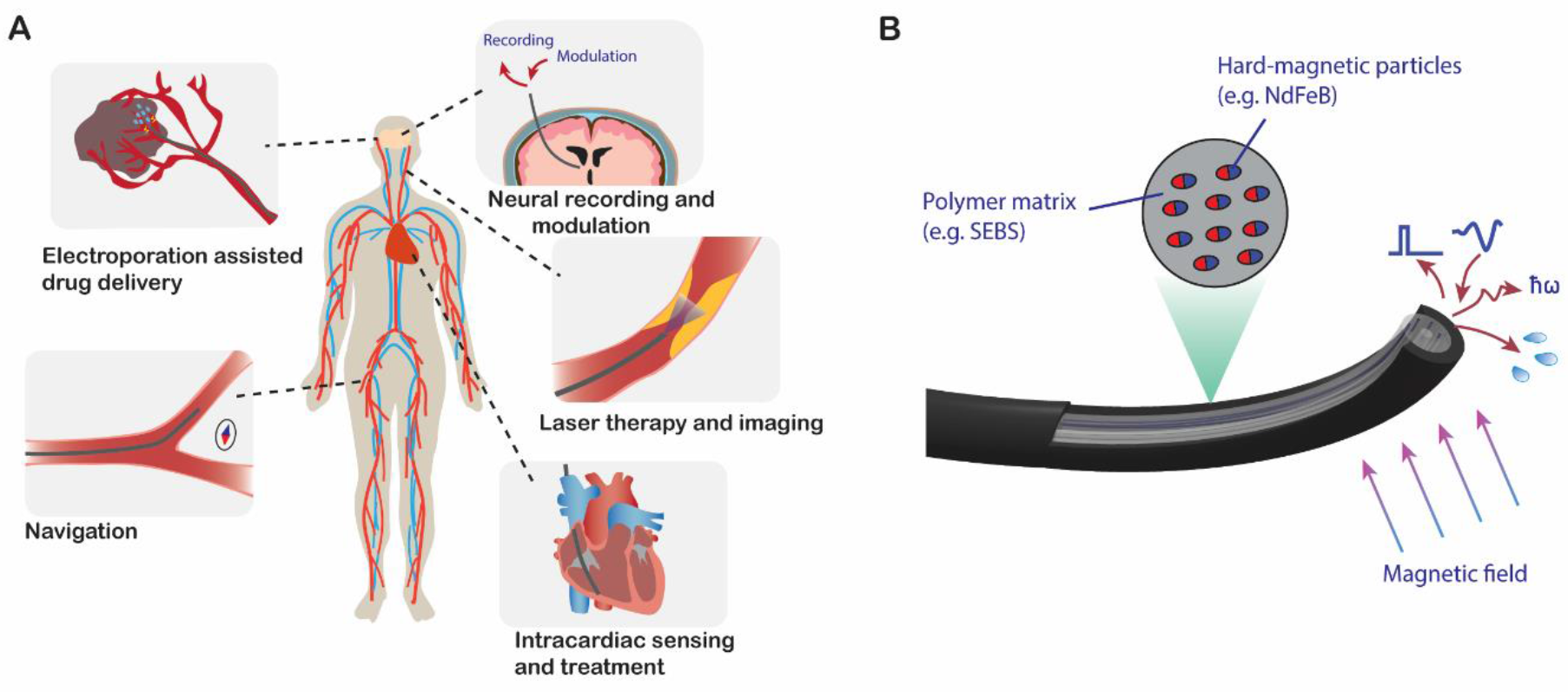
Multifunctional ferromagnetic fiber robots (MFFRs) for minimallyinvasive surgery. (**A**) Clinical procedures that MFFRs can be used for, including navigation inside blood vessels, electroporation-assisted drug delivery to overcome blood-brain barriers, deep brain neural recording and modulation, laser therapy for removing blood clots and endovascular imaging, and intracardiac sensing and treatment. (**B**) Schematic illustration of fiber robots that are composed of soft polymer matrix with dispersed hard-magnetic particles and a functional core. The magnetic polarity is along the fiber axial direction. The functional core consists of waveguides, electrodes, and microfluidic channels that can deliver and record optical and electrical signals and transport liquid.

## Results

### Design and fabrication of multifunctional ferromagnetic fiber robots

In this work, multiple MFFRs structures were designed and developed for various application scenarios. The MFFRs consist of thermoplastic elastomer jackets doped with hard-magnetic particles and multifunctional cores with various components, such as waveguides, microfluidic channels, and electrodes (**Fig. 1B**). The integrated materials and components allow for magnetically controlled steering, optical signal delivery and collection, fluid delivery, and electrical stimulation and recording. Besides, this functional core also provides the mechanical support required for probe insertion during surgeries.

To fabricate the MFFRs, we use a scalable thermal drawing process, in which a macroscopic “preform” with predesigned structures is thermally drawn into a microscale fiber with preserved cross-sectional geometries. **Supplementary Fig. 1** shows a representative “preform” preparation process to create an MFFR, which consists of a core with a waveguide, an electrode, and a hollow channel, a ferromagnetic middle layer, and a sacrificial outer layer as a support for the thermal drawing process. The ferromagnetic component (FC) was prepared by evenly dispersing hard-magnetic microparticles (neodymium iron boron, NdFeB) in thermoplastic elastomers using a hot press (see Method for details). We chose Styrene-ethylene-butylene-styrene (SEBS) as the thermoplastic elastomer material owing to its low elastic modulus, good biocompatibility, and compatibility with the TDP^29,36^. In this work, we used multiple types of electrodes, including low-melting-point metals (e.g., BiSn) and high-melting-point metals (e.g., Ag). Waveguides used include step-index polymer waveguides made of polycarbonate (n = 1.58) core and polymethyl methacrylate (PMMA, n = 1.49) cladding, commercially available PMMA waveguides, and silica waveguides.

The prepared macroscopic preform was then heated above the glass-transition temperature of the polymers and pulled into a ~150-m-long fiber (**Supplementary Fig. 2**) at a speed of about 4 m/min under an applied high temperature (260-300 °C) and external stress (**Fig. 2A**). The drawn fiber has the same cross-sectional geometry and composition as the “preform” but with a 20-150-fold reduction in dimensions. We employed a loading concentration of 35% (v/v), which is the highest concentration that we could achieve when drawing at this temperature. With a higher loading concentration, the viscosity of the composite becomes too high, making it not suitable for the thermal drawing process. Using this method, we fabricated fiber F1 (**Fig. 2B**) with one polymer waveguide, one BiSn electrode, and one microfluidic channel, and fiber F2 (**Fig. 2C**) with one polymer waveguide and four hollow channels. Alternatively, materials with high melting temperatures, such as silver wire and silica waveguide, can also be integrated into the MFFR via a convergence drawing process (**Fig. 2D**). Silver wires or silica waveguide were threaded through the channels inside the preform and converged with the surrounding polymer during the pulling down procedure, resulting in fiber F3 (**Fig. 2E**) with two silver electrodes and one microfluidic channel, and fiber F4 (**Fig. 2F**) with one silica waveguide and four microfluidic channels. After thermal drawing, the fiber tips (≈2.5 cm long segments) were magnetized along the fiber axis with a magnetic field of 2.2 T using a high-field electromagnet. It is worth noting that the high temperature applied during the drawing process had only a modest effect on the magnetic properties of the NdFeB composite. The magnetization loop of the NdFeB composite (20 % (v/v)) as shown in **Fig. 2G** indicates that the magnetization became saturated when the applied magnetic field strength reached 2 T. After the drawing process, the remanent magnetization of the composite dropped by only < 10%, changing from 116±2 kA/m to 109±2 kA/m (**Fig. 2H**, n=4).

**Fig. 2.**
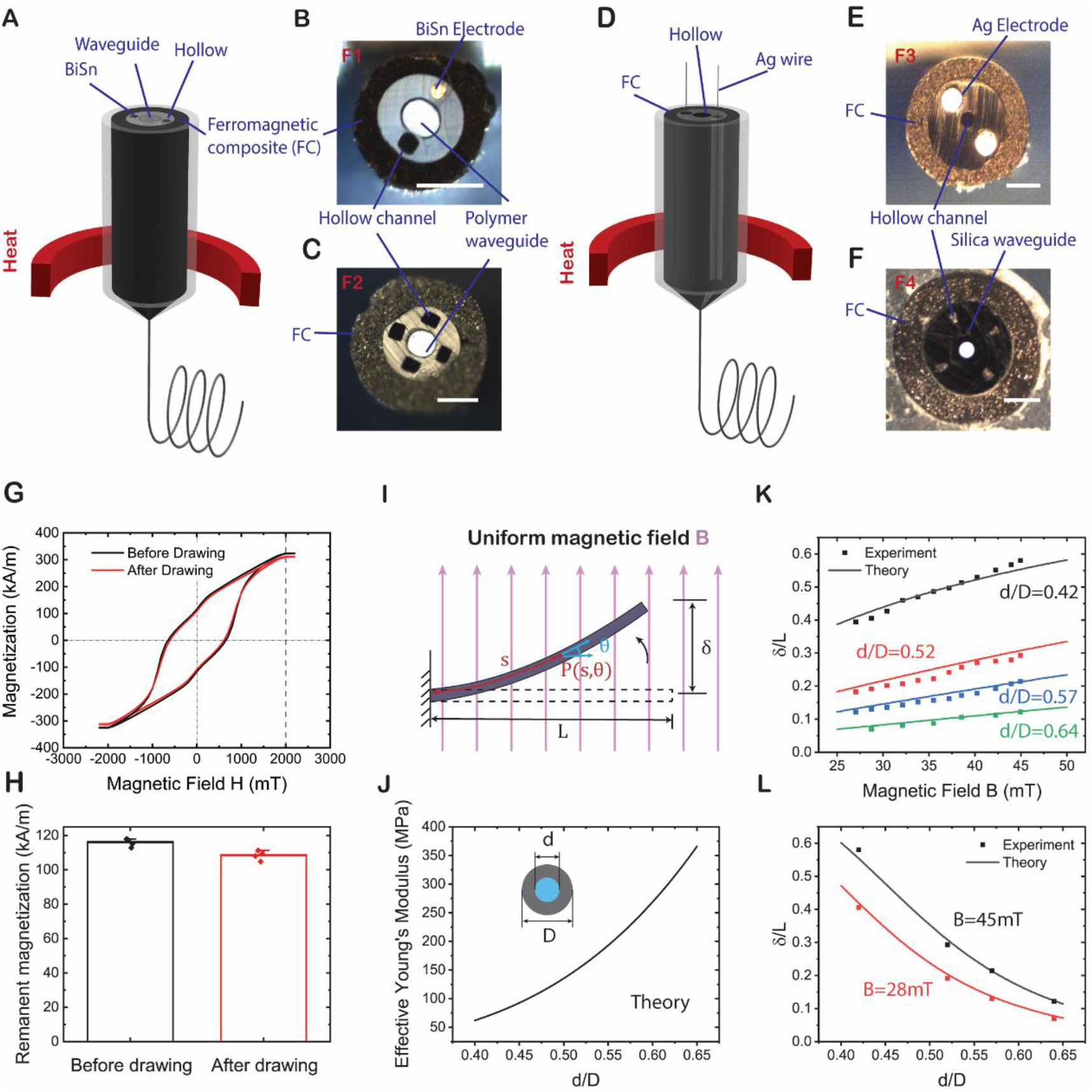
Design and fabrication of MFFRs. (**A**) A schematic of the thermal drawing process and the drawn fiber with a jacket made of ferromagnetic composite (FC) and a functional core consisting of (**B**) a polymer waveguide, a BiSn electrode, and a hollow channel, and (**C**) a polymer waveguide and four hollow channels. (**D**) A schematic of the thermal convergence drawing process and the drawn fiber with a jacket made of FC and a functional core consisting of (**E**) a pair of Ag electrodes and a hollow channel, and (**F**) a silica waveguide and four hollow channels. (**G**) Magnetization loops of the hard-magnetic composite before and after the thermal drawing process. (**H**) Comparison of the remanent magnetization of the hard-magnetic composite before and after the thermal drawing process, indicating a <10% reduction of the magnetization strength. (**I**) Schematic of a ferromagnetic fiber robot deflecting toward the direction of the uniform magnetic field **B** applied perpendicularly to the fiber. The unconstrained length of the robot is denoted L. δ indicates the deflection of the free end. *s* denotes the arc length from the fixed point to the point of interest (denoted by *P*), *θ* denotes the angle between the tangent to the curve at point *P* and the reference direction. (**J**) Calculated effective Young’s modulus of the fiber robots plotted against core-to-fiber ratio. The fiber structure is simplified as a polycarbonate core (diameter, d) and an FC jacket (outer diameter, D, loading fraction, 35 %(v/v)). (**K**) Normalized deflection δ/L predicted from theory and experimental measurements plotted against the applied field strength with different core-to-fiber ratios: d/D = 0.42, 0.52 0.57, 0.64 when *L/D* = 40. (**L**) Normalized deflection δ/L predicted from theory and experimental measurements plotted against core-to-fiber ratios under different magnetic field strengths: B = 28 mT, 45 mT. Scale bar, 150 μm. All error bars and shaded areas in the figure represent the standard deviation.

The magnetic actuation property of the MFFRs mainly depends on their magnetic and mechanical properties, both of which are affected by the fiber geometry and particle loading concentration. Here, we explored the effects of these factors analytically by using a simplified model, where the fiber was considered as a beam with a length *l* and placed perpendicular to a uniform magnetic field (**Fig. 2I**). The diameters of the inner functional core and the overall fiber are *d* and *D*, respectively. The governing equation describing the fiber response to a uniform magnetic field *B* can be expressed as^20^

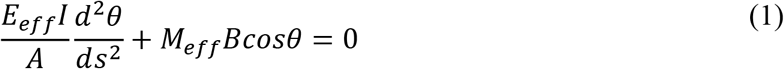

where *s* denotes the arc length from the fixed point to the point of interest (denoted by *P* in **Fig. 2I**), *θ* denotes the angle between the tangent to the curve at point *P* and the reference direction, *E*_*eff*_ is the effective Young’s modulus of the whole beam, *I* is the area moment of inertia, which can be expressed as *I* = *πD*^4^/64 for a cylindrical beam, *A* is the cross-section area of the whole beam, which can be expressed as *A* = *πD*^2^/4, and *M*_*eff*_ is the effective magnetization density of the whole fiber, which can be described as

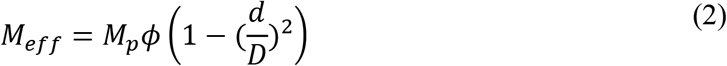

where *M*_*p*_ denotes the magnetization density of the particles, and *ϕ* denotes the particle loading volume concentration. The effective Young’s modulus of the fiber can be calculated by^21^

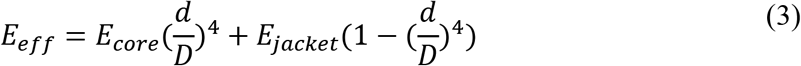

where *E*_*core*_ and *E*_*jacket*_ denote the Young’s moduli of the fiber core and ferromagnetic microparticles loaded SEBS. To simplify the model, we use a fiber structure that has a polycarbonate core without features and set *E*_*core*_ as 2.4 GPa. **Fig. 2J** shows that the effective Young’s modulus of the fiber increases nonlinearly as the core-to-fiber ratio *d/D* increases when *E*_*jacket*_ is set as 11 MPa with a particle doping concentration of 35 %(v/v). We choose a *d/D* range of 0.4 to 0.65 for further investigation. When *d/D* is too small, the fiber becomes too soft, and thus cannot provide the stiffness and mechanical strength required for the probe insertion process. On the other hand, a larger *d/D* results in a larger stiffness of the fiber, which limits its actuation response under a magnetic field.

By substituting Eqs. 2 and 3 into Eq. 1, we can obtain the deflection of the magnetically active fiber tip (details are available in the Supplementary Materials). The predicted fiber deflection ratios *δ*/*L* from modeling show good agreement with experimental data (**Fig. 2K** and **2L**) under various core-to-fiber ratios *d/D* and magnetic field strengths. The normalized deflection ratios *δ*/*L* of the fibers are 0.12, 0.21, 0.31, and 0.58 at 45 mT for different core-to-fiber ratios: *d/D* = 0.42, 0.52 0.57, and 0.64 when *L/D* = 40, respectively. When other conditions remain unchanged, a smaller core-to-fiber ratio will lead to a smaller effective Young’s modulus and a larger effective magnetization density, resulting in a larger response to the same external magnetic field. Also, when the magnetic particles loading concentration *ϕ* and core-to-fiber ratio *d/D* are in the range of 0 −35 %(v/v) and 0.4 −0.7, respectively, the fiber deflections increase with the loading concentration (Supplementary Fig. 2). Based on this model, we can design fiber robots by tuning their geometries and materials to fit the mechanical and magnetic actuation requirements for different application scenarios.

### In vitro evaluation of the MFFRs in 2D and 3D vascular phantoms

Hereafter, we demonstrated MFFR’ magnetic steering capability, as well as additional functionalities enabled by the functional core using 2D and 3D vascular phantoms. The 2D phantom was designed to have multiple bifurcations and branches with different angles, with a channel width of 2.5 mm. First, we performed the navigation using a commercial J-tip guidewire (diameter, 0.36 mm, Supplementary Fig. 4) with an angled tip (**Fig. 3A** and **Movie S1**). Steered solely through manual rotation of its proximal end, the guidewire could get into the branch at an angle of 60° or below. However, the guidewire was unable to pass the bifurcation with an angle of 70° after repeated attempts. Note that this guidewire didn’t have any additional sensing or treatment functionalities. **Fig. 3B** and **Movie S2** show a representative case achieved through a fiber robot (same structure as F2, diameter, 0.38 mm, **Supplementary Fig. 4**) under the guidance of an external magnetic field. During the demonstration, the fiber robot was manually fed and retracted while the tip was remotely controlled by magnetic fields generated by a permanent magnet (diameter and height of 50 mm). The typical working distance between the fiber tip and the magnet ranged from 30 to 60 mm, which corresponds to the field strength ranging from about 40 to 150 mT. The fiber robot could smoothly navigate through the targeted path by passing three bifurcations with angles of 70°, 50°, and 30° without any noticeable difficulties. Although the demonstration of a certain angled guidewire could not represent the guidewire manipulation performed by professional interventional physicians, it was evident that the use of the proposed fiber robots could improve the navigation capabilities via magnetic steering, especially in tortuous vascular environments where bifurcations had highly angled branches.

**Fig. 3.**
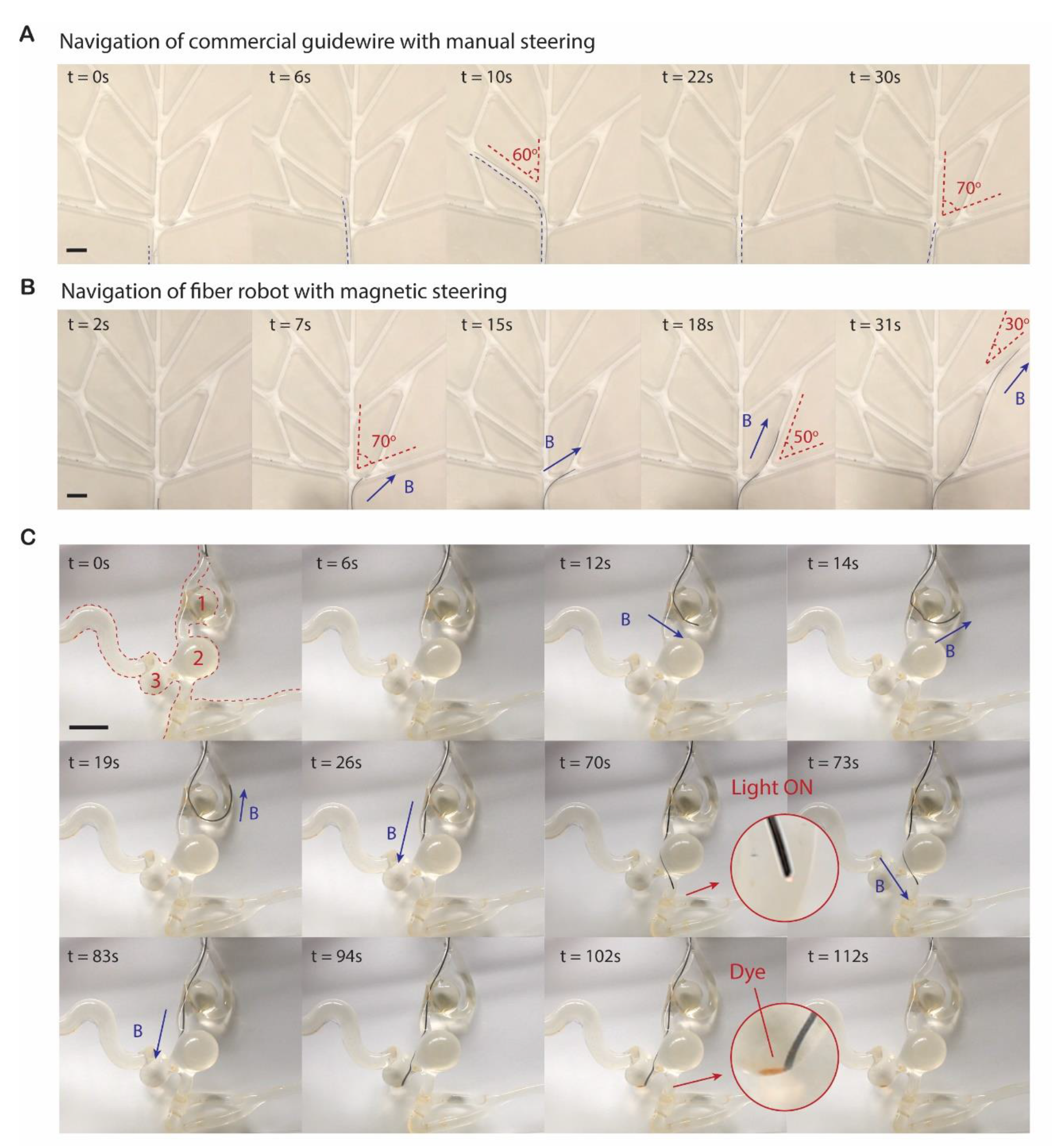
In vitro evaluation of the MFFRs in 2D and 3D vascular phantoms. (**A**) Navigation of a commercial guidewire (diameter, 0.36 mm) with manual steering in a 2D vascular phantom model with multiple bifurcations. The diameter of the channels was 2.5 mm. Blue dashed lines indicate the direction of the guidewire. Red dashed lines indicate the bifurcation angles. The guidewire in the picture may appear thinner than the actual size due to the low contrast with the background. (**B**) Navigation of the fiber robot (same structure as fiber F2, diameter, 0.38 mm) with magnetic steering in a 2D vascular phantom model. (**C**) Navigation and operation of the fiber robot (fiber F2, diameter, 0.5 mm, microfluidic channel dimensions, 65 × 50 μm) in a 3D vascular phantom model. Orange dashed lines show the outline of the model. The fiber robot made a sharp turn to the vessel near the first aneurysm (t = 19s) with magnetic steering, delivered light (t = 70s) after passing the second aneurysm, and injected red dye (t = 102s) into the third aneurysm. An external magnetic field (40 to 95 mT) was generated by a cylindrical (diameter and height of 50 mm) permanent magnet at a distance (40 to 60 mm). The proximal end was manually pushed to advance the fiber robot during the navigation. The fiber robot in the pictures may appear thicker than the actual size due to the magnifying effect of the round wall of the vessel phantom. Scale bar, 10 mm.

We further performed another demonstration in a real-sized, 3D silicone human vascular phantom, with three aneurysms (**Fig. 3C** and **Movie S3**). The inner diameter of the silicone vessels along the targeted path ranged from 2.5 to 3.5 mm, while the diameters of the three aneurysms were 12 mm (first), 14 mm (second), and 10 mm(third). Thanks to the functional core, we could achieve not only magnetic navigation, but also other potential therapeutic functions, such as light therapy and fluid delivery, using a single fiber robot. Light and optical technologies are widely used in modern medicine for diagnosis, therapy, and surgery^37^, including laser angioplasty, photodynamic therapy, photothermal therapy, and light-triggered drug release. Also, conventional catheters often contain tubes that enable localized delivery of contrast agents and embolization agents. To validate that our fiber robots can potentially achieve these functionalities, we used fiber F2 (diameter, 0.5mm, microfluidic channel dimensions, 65 × 50 μm) with one waveguide, four microfluidic, and an FC jacket for the demonstration. Under the guidance of an external magnetic field, the fiber robot reached the vessel near the first aneurysm with a sharp turn at t = 19s. After that, it changed the direction, passed the second aneurysm, and emitted red laser light. Next, it turned to another branch, and injected red liquid dye into the third aneurysm. These results demonstrated that the MFFRs enable remote steering with larger angles compared with conventional guidewires owing to its active steering capability, and simultaneous therapeutic treatment including light delivery and drug injection.

### Endocardial electrogram (EGM), pacing, and bioimpedance monitoring in a Langendorff-perfused mouse heart model

Next we demonstrated the application of the fiber robots in interventional endocardia surgery using a human cardiac model (**Fig. 4A** and **Movie S4**). The fiber robot (F3) was inserted from the superior vena cava, while the red dashed line indicates the original direction. Under the guidance of an external magnetic field (20 to 60 mT) generated by a permanent magnet at a distance (50 to 70 mm), the fiber reached the target site at the right ventricle near the apex. Then we injected red dye from the microfluidic channel inside the fiber to indicate the location of the fiber tip. To further validate the diagnostic and therapeutic functions of the multifunctional fiber robots, we performed EGM, pacing, and bioimpedance monitoring experiments using a Langendorff-perfused mouse heart model. To mimic the scenario in interventional cardiovascular surgeries, we inserted fiber F3 from the superior vena cava of the mouse heart. The fiber passed the right atrium and tricuspid valve, and finally reached the right ventricle with the tip gently touching the inner wall close to the apex (**Fig. 4B**). The whole heart electrocardiogram (ECG) was recorded with three leads in the bath while localized bipolar EGM was recorded simultaneously through two exposed silver electrodes at the fiber tip, implying good contact between the fiber and the ventricle wall (**Fig. 4C**). The bipolar EGM reached the negative peak when the cardiomyocytes underneath the electrodes depolarized^38^. This depolarization happened between the R peak (apex ventricle activation) and S peak (base ventricle activation) recorded in ECG, which also verified that the fiber tip was localized between the apex and base of the right ventricle. Next, we delivered electrical pulses through the two electrodes inside the fiber and successfully paced the heart as shown in the recorded ECG (**Fig. 4D**). Before pacing, the R-R duration was around 220 ms. Upon pacing, the heart was forced to follow the paces of the stimulating pulses with a cycle length of 150 ms. After pacing for about 20 s, we stopped the stimulating signal, and the heart gradually recovered to the original stage with an R-R duration of ~ 220 ms.

**Fig. 4.**
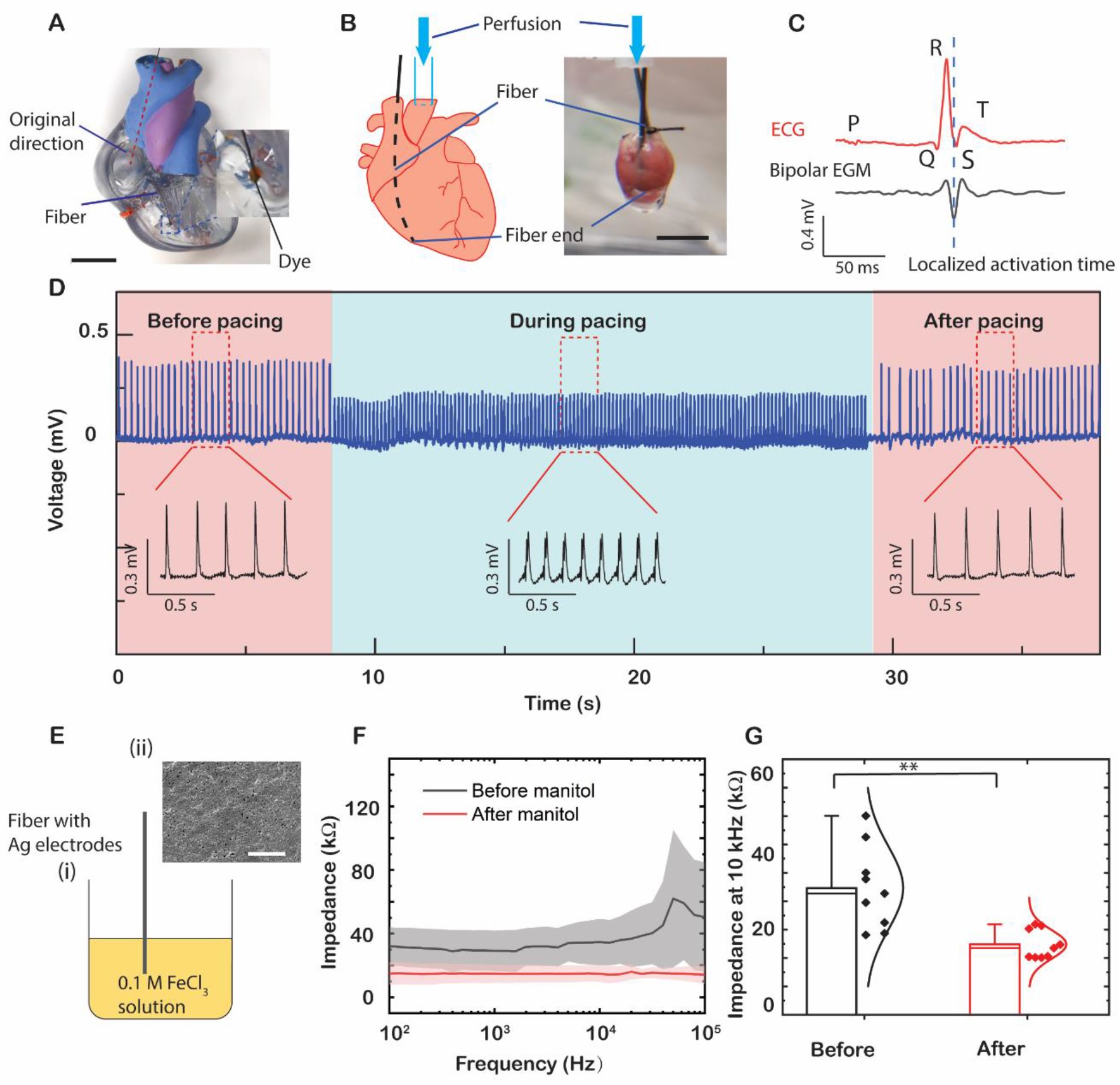
Endocardial electrogram (EGM), pacing, and bioimpedance monitoring in Langendorff-perfused mouse heart model. (**A**) A demonstration of the application of the fiber robot during interventional endocardial surgery using a human heart model. The fiber robot (fiber F3, diameter, 0.45 mm) was magnetically guided to the targeted site in the right ventricle. Red dye was injected from the microfluidic channel inside the fiber. An external magnetic field (20 to 60 mT) was generated by a cylindrical (diameter and height of 50 mm) permanent magnet at a distance (50 to 70 mm). The red dashed line indicates the original direction of the fiber without magnetic field. Scale bar, 30 mm. (**B**) Schematic and photograph of the test setup. The fiber (F3) was inserted from the superior vena cava, passed the right atrium and tricuspid valve, and finally reached the inner wall of the right ventricle. Scale bar, 5mm. (**C**) A representative whole heart ECG trace and localized bipolar EGM trace recorded simultaneously through leads in the bath and two exposed electrodes at the fiber tip, respectively. (**D**) ECG of the mouse heart recorded before, during, and after pacing with a cycle length of 150 ms. (**E**) Surface chlorination of Ag electrodes by immersing the fiber into FeCl3 solution (i) and SEM image of the chlorinated surface (ii). Scale bar, 5 μm. (**F**) Impedance spectra of the mouse heart measured using the fiber before and after mannitol perfusion. (**G**) Comparison of the mouse heart impedance at 10 kHz before and after mannitol perfusion. (Welch’s *t*-test, *p*-value: 0.003, \*\**p*<0.01, n=9 measurements, n=3 hearts) All error bars and shaded areas in the figure represent the standard deviation.

Myocardial tissue impedance decreases with cardiac edema which is associated with heart disease^39,40^. Thus, it is important to monitor myocardial tissue impedance during cardiovascular surgeries, such as off-pump coronary artery bypass (OPCAB) surgery^41^. In order to investigate if our fiber could monitor myocardial tissue impedance in the presence of cardiac edema, we performed two-electrode bioimpedance measurements through electrodes inside the fiber before and after mannitol perfusion. To obtain a lower noise level and better chemical stability, we chlorinated the surface of the silver electrodes by immersing the fiber in FeCl_3_ solution for 1 min (**Fig. 4E(i)**). The scanning electron microscopy (SEM) image (**Fig. 4E(ii)**) shows the sub-micron structures on the tip surface after chlorination, and the corresponding energy-dispersive X-ray spectroscopy (EDS, **Supplementary Fig. 5**) verifies the presence of the Ag/AgCl layer. From the measured impedance spectra, we observed that the myocardial impedance dropped significantly after mannitol perfusion (**Fig. 4F**), while the perfusion solution impedance stayed consistent (**Supplementary Fig. 6**). Specifically, at 10 kHz, the myocardial impedance dropped from 35±15 kΩ to 15±5 kΩ (**Fig. 4G**, Welch’s *t*-test, *p*-value: 0.003, n=9 measurements, n=3 hearts). We believe this impedance drop is due to mannitol-induced extracellular edema which was reported previously^42,43^.

**Fig. 5.**
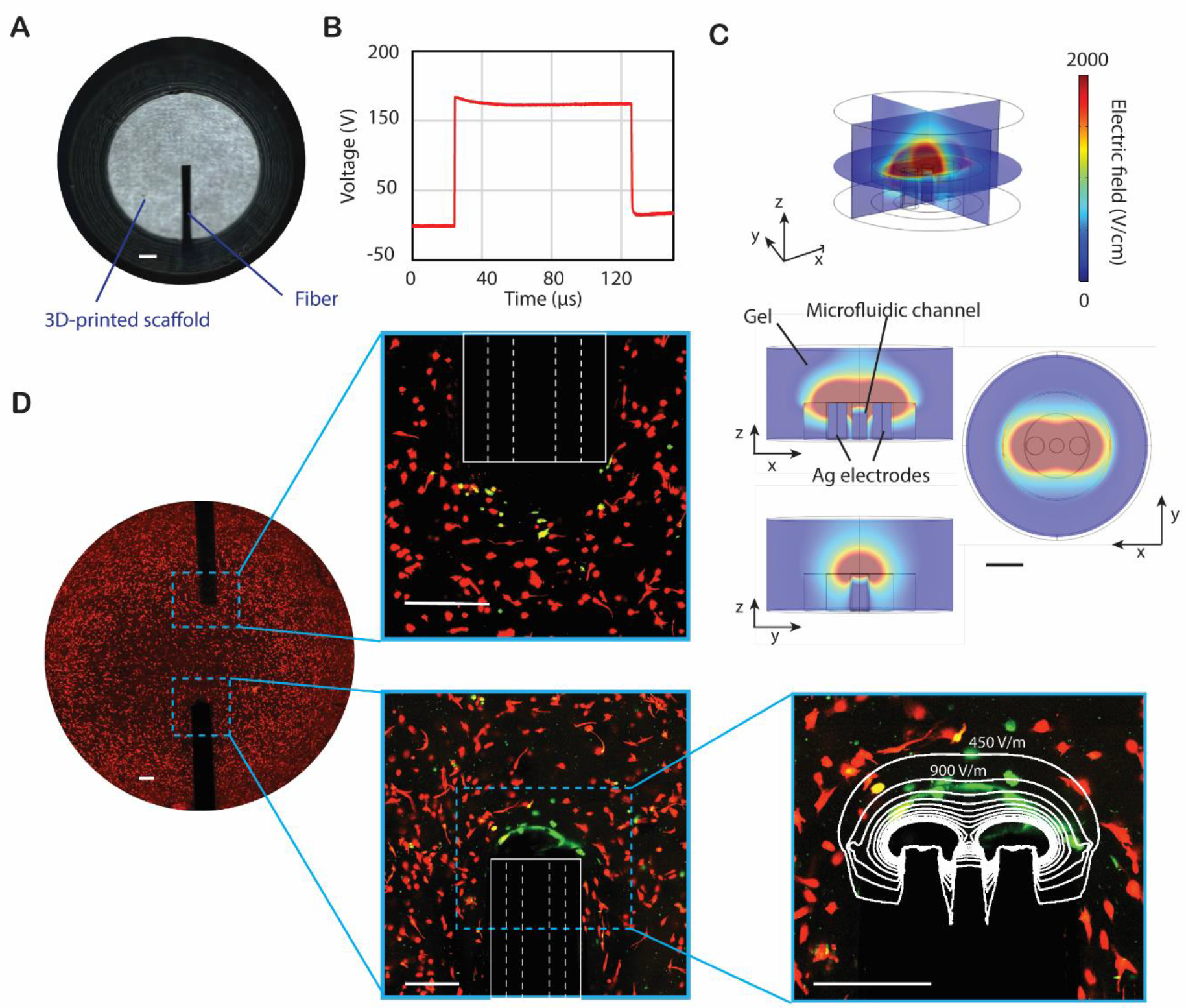
Simultaneous microscale electroporation and chemical delivery. (**A**) Experiment setup with fiber robots in a 3D-printed scaffold. Scale bar, 1mm. (**B**) A typical voltage waveform (175 V, 100 μs pulse width). (**C**) FEA results of the 3D electric field distribution during electroporation. The positions of the Ag electrodes, microfluidic channel, and gel are indicated in the middle frame. Scale bar, 200 μm. (**D**) A representative image of the reversibly electroporated cells indicated by the YO-PRO-1 (green) surrounded by unaffected cells indicated by Calcein red AM (red). The right frames are zoomed-in images of the blue dashed boxes on the left. The white boxes in the middle frames indicate the fiber robots while the white dashed lines indicate the embedded electrodes. The white lines in the bottom right frame indicate the corresponding contour plot of electrical field intensity obtained from FEA. Scale bar, 300 μm.

Compared to conventional millimeter scale catheters, our fiber robots with a submillimeter size and active steering capability offer extra benefits in precise control and localized sensing, especially for operation on small hearts. For instance, cardiac intervention for pediatric patients with congenital heart disease requires small devices for both diagnostic and therapeutic catheterization^44,45^. Pre-clinical cardiac research is another potential application of our fiber robots. The current electrophysiological research on isolated heart models is limited to the measurement of epicardial action potential^46,47^ and bioimpedance^48^. Our miniaturized fiber robots can provide sensing and modulation tools for cardiac research involving endocardial measurement in the intact small heart. With the platform we developed, these fiber robots can potentially be improved to sense more physiological conditions, including temperature, blood pressure, blood oxygen, etc.

### Simultaneous microscale electroporation and chemical delivery

Electroporation is the phenomenon of creating pores in the lipid bilayers of cell membranes when the applied potential differences are greater than the transmembrane voltage. These pores can be either reversible or irreversible, depending on their size and the number of pores formed. Specifically, reversible electroporation can be combined with drug delivery to treat disease or infection sites and increase the synergistic uptake of drugs.

To evaluate the electroporation and drug delivery capability of the fiber robots, we cultured U-251 glioblastoma cells in hydrogels in 3D-printed scaffolds (**Fig. 5A**) and delivered reversible electroporation (RE) treatments to produce a volume of electroporated cells. Glioblastomas are the most commonly occurring cranial tumor, with a life expectancy under current treatment modalities averaging a little over a year after diagnosis^49^. U-251 cells are between 20 to 40 μm in length and have an average area of 700 μm^2^ in a 5 mg/mL collagen hydrogel. We inserted a fiber with two Ag electrodes and one microfluidic channel (fiber F3 in **Fig. 1E**) into the gel. During the testing, electric pulses were applied through the two electrodes while 5mM of Ca^2+^ adjuvant was introduced through the microfluidic channel to reduce the electric field threshold (EFT). Pulse parameters were set to 200 pulses with a pulse width of 100 μs, pulse amplitude of 175 V, and a repetition rate of 1 Hz **(Fig. 5B**). To simulate the electric field distribution during electroporation, we also performed finite element analysis (FEA) of the experiment, as shown in **Fig. 5C**. Due to the minimal gap of 140 μm between the charged leads and the insulated walls of the conducting wires, the electric field decays rapidly from the fiber surface. Simulations also showed no observable temperature increase over the entire treatment. **Fig. 5D** shows the *in vitro* reversible electroporation results. Unaffected cells express red fluorescence while membrane-electroporated cells express green fluorescence due to the uptake of YO-PRO-1 molecules. The measured RE EFT was between 450-500 V/cm, higher than that reported in the literature^50^. This may be due to the small number of cells present within the lesion, reducing the accuracy of lesion size. The negligible heating of the gel may be another reason, as temperature rise is known to reduce EFTs.

These results show that we can achieve effective localized reversible electroporation treatment and chemical delivery simultaneously through the fiber robots.

### In vivo neural electrophysiological recording and optogenetic control

Implantable neural probes that penetrate brain tissue can record various types of electrophysiological signals and perform neural modulation^51^. However, it is challenging to target the probes to the desired location precisely, especially under complex conditions with obstacles, such as significant fiber tracts, tumors and vessels^52^. These complications can be significantly minimized by using steerable and flexible devices, such as magnetic needles^53,54^, which can reach deep brain region using curved trajectories and bypass critical structures. Here, with the method we developed, we miniaturize the size of the device and the produce a steerable multifunctional neural probe with an overall diameter of 0.3 mm (fiber F1 in **Fig. 2B**). With the magnetically steerable jackets and the assistance of external magnetic fields, we can insert the neural probes in a more controllable and precise manner by avoiding obstacles, allowing access to some out-of-reach places, and reducing neurovascular damages (**Fig. 6A**). As shown in **Fig. 6B**, without an external magnetic field, the probe was straightly implanted in a brain phantom (0.6 wt% agarose gel). With the control of a permanent magnet, the probe can be implanted with a deflected angle of about 45°. We then injected red dye through the inner microfluidic channel to indicate the location of the probe tip. Besides, the impedance of the fiber probes at 1kHz was found to be 271±46 KΩ, making them suitable for spike recording (**Fig. 6C**).

**Fig. 6.**
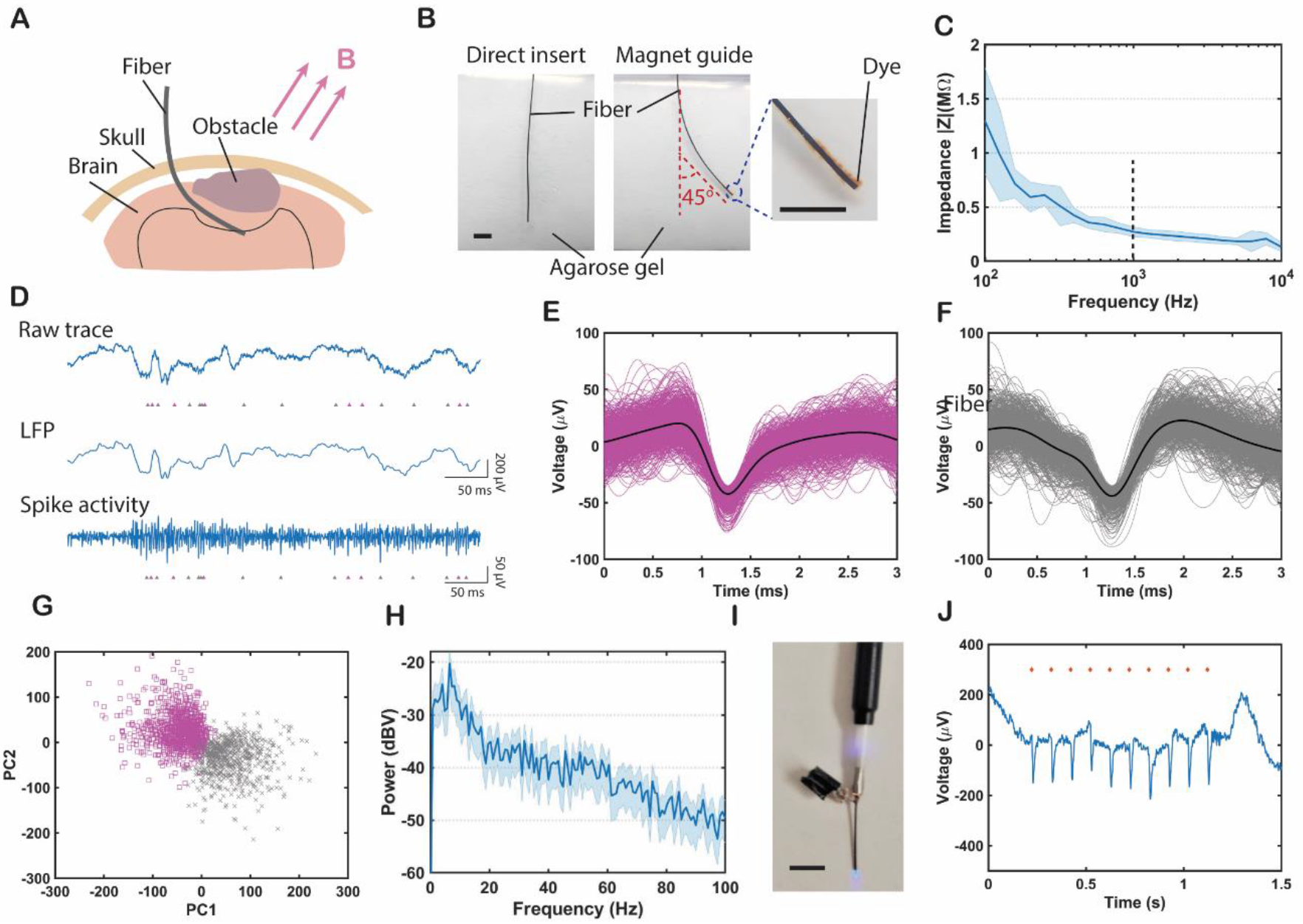
In vivo neural electrophysiological recording and optogenetic control. (**A**) A schematic showing the ferromagnetic fiber robots bypassing obstacles during neural probe insertion. (**B**) Validation of the magnetically steerable fiber robots (fiber F1, diameter, 0.3 mm) in the brain phantom (0.6% agarose gel). The probe was deflected by an angle of about 45° during insertion with the magnetic guidance. An external magnetic field (60 to 200 mT) was generated by a cylindrical (diameter and height of 50 mm) permanent magnet at a distance (25 to 50 mm). Red dye was injected from the microfluidic channel inside the fiber. Scale bar, 4 mm. (**C**) Impedance measurement of the BiSn electrode in fiber F1. (**D**) Recorded electrophysiological signal via fiber F1 in the hippocampal region of mice brain. The top trace shows the unfiltered signal, the middle trace shows bandpass-filtered (0.3–300 Hz) LFP, and the bottom trace represents the bandpass-filtered (0.3–5 kHz) spike trace. Pink and grey markers indicate the time points of two different units. (**E**, **F**) Action-potential shapes of the two units. (**G**) Principal-component analysis (PCA) of the two units. (**H**) Power spectral density analysis of the LFP. (**I**) A photograph of the assembled device coupled to the 473 nm laser. Scale bar, 5mm. (**J**) Unfiltered electrophysiological recording from fiber F1 during optogenetic stimulation (10 Hz, 5ms pulse width, 5.1 mW/mm^2^). (n=4) All shaded areas in the figure represent the standard deviation.

To evaluate the functionality of the fiber probes in recording localized brain activities, we implanted the fiber probes into the hippocampal region of wild-type mice (n=4). A representative electrophysiology recording including raw trace, local field potential (LFP, 0.3 to 300 Hz), and spike activities (0.3 to 5 kHz) is shown in **Fig. 6D**. The embedded electrode captured spiking activities from multiple neurons, from which we found two distinctly insulated clusters via principal component analysis (PCA) (**Fig. 6E-G**). The quality of the isolation was evaluated by L-ratios and isolation distance, which were 0.05 and 75, respectively. The two clusters were also observed in the raw trace and spiking traces as marked with the corresponding matched colors in PCA, which demonstrates the fiber probes can record neural activities with a single-unit resolution. Besides, the power spectral density analysis of the recorded trace in **Fig. 6H** shows that brain oscillations were observed in the frequency range of 6-10 Hz, corresponding to the theta oscillations in the hippocampal network pattern of activity in mice^55^.

Furthermore, we demonstrate that simultaneous optical modulation and electrophysiological readout can be achieved using the multifunctional fiber. Transmission spectroscopy confirmed the utility of the probes for optical guidance in the visible range (450–750 nm) (**Supplementary Fig. 7A**). Specifically, the transmission attenuation measured using the cut-back method is 0.797 dB/cm at a wavelength of 473 nm (**Supplementary Fig. 7B**), which is the excitation peak of channelrhodopsin2 (ChR2). During the experiment, the waveguide inside the fiber was coupled to a silica optical fiber using a direct ferrule-to-ferrule coupling, while the electrode was electrically connected to a pin (**Fig. 6I**). Then the fiber probes were implanted in Thy1-ChR2-YFP mice, which express ChR2 across the nervous system, in the hippocampal region (n=4). We applied laser pulses at a frequency of 10 Hz with a pulse width of 5 ms and a power density of 5.1 mW/mm^2^ through the fiber probes, and recorded optically evoked neural electrophysiological activities using the electrodes inside the fiber probes, as shown in **Fig. 6J**. These results demonstrate that the multifunctional fiber robots can be magnetically steered in a brain phantom, and can record single-unit electrophysiological signals, and perform optogenetic stimulation and recording in mice.

### Fiber bundle endoscope

Optical fiber bundle endoscopes are widely used for imaging, sensing, and illumination in hard-to-reach locations of the human body. With the technology that we developed, we can produce fiber bundle endoscopes with active steering in a single step. The fiber here consisted of a fiber bundle core for imaging and a ferromagnetic jacket for steering (inset figure in **Fig. 7A**). PMMA fibers were chosen for imaging pixels due to their flexibility and low melting temperature. The diameter of the imaging core was about 350 *μm* with 320 pixels. The size of each pixel was about 20 μm and the overall diameter of the fiber was about 600 μm. The number of pixels and imaging resolution can be further improved by using smaller fibers for bundling in the preform and increasing the draw-down-ratio during the TDP. The transmission spectrum shows that this thermally drawn imaging fiber could guide light across the visible range (**Supplementary Fig. 7C**) and the attenuation is ~0.316 dB/cm at a wavelength of 615 nm (**Supplementary Fig. 7D**). **Fig. 7A** shows the experimental setup for imaging demonstration. For our object generation, we used a halogen broadband light that is delivered by a multimode fiber, and the light was focused onto a custom-made mask. Then the pattern on the mask was imaged onto the distal end of the imaging fiber. During the experiment, the pattern focused on the fiber end was enlarged by 1.2 times compared to the original mask (**Fig. 7B**) after the lens set. Images (**Fig. 7C**) were collected from the other end of the fiber through a camera. For demonstration, two different patterns (a 100-μm-thick line and a character “C”) were successfully captured via a 40-cm-long imaging fiber with distinguishable images.

**Fig. 7.**
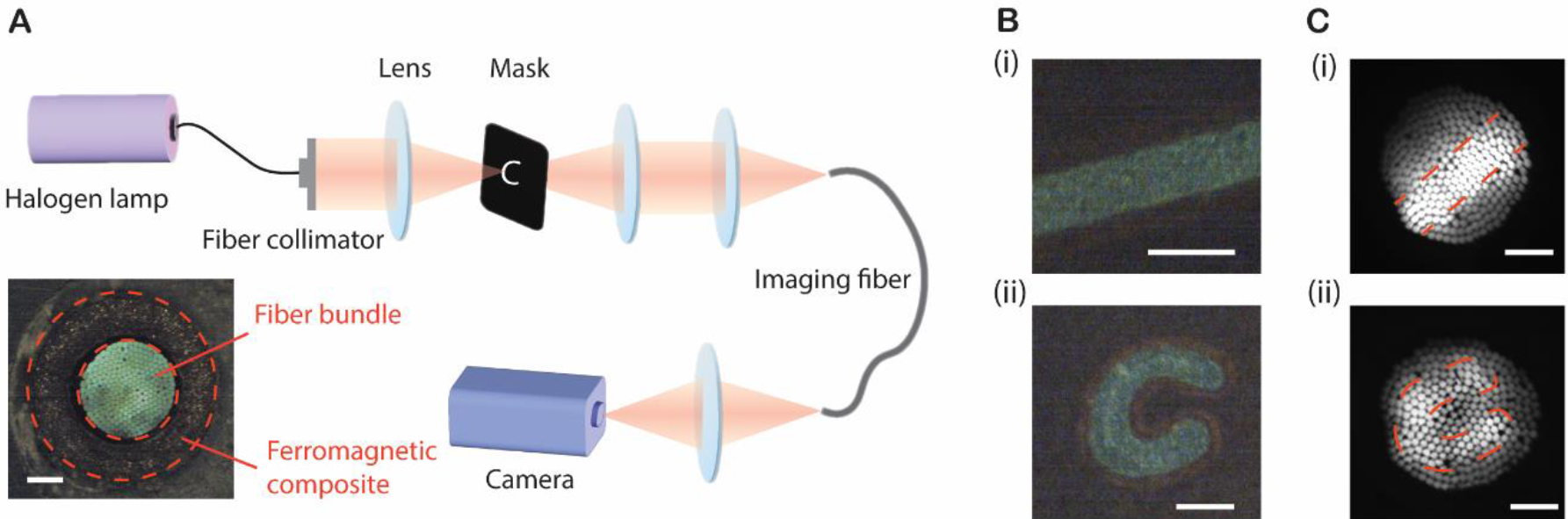
Fiber bundle imaging. (**A**) A schematic of the experimental setup for imaging demonstration and (inset) the cross-section image of a thermally drawn imaging fiber with an imaging core and a ferromagnetic jacket. The pattern on the mask was imaged onto the tip of the imaging fiber after the lens sets with a magnification of 1.2. This image was transmitted to the other end of the fiber, from where it was relayed onto a CCD camera. The pixel size of the imaging fiber is about 20 μm and the pixel count is 320. Scale bar of the inset, 150 μm. (**B**) shows the patterns on the mask and (**C**) shows the corresponding patterns captured via a 40-cm-long imaging fiber. Scale bar, 100 μm.

## Discussion

The results presented here address several limitations of conventional robotic devices through a variety of miniaturized MFFRs fabricated using the thermal drawing process. The thermal drawing process allows the scalable manufacturing of multimaterial fiber robots with the integration of ferromagnetic, electrical, optical, and microfluidic components. Each drawing can produce fibers of more than 100 meters with sizes as small as 250 μm at a speed of about 4 m/min. These MFFRs can steer through complex and constrained environments with a controlled external magnetic field due to the embedded ferromagnetic microparticles. In order to significantly reduce the size of the fiber robots, we abandon the conventional guidewire/catheter system and combine navigation, diagnosis, and therapeutic functions in a single fiber device. These multifunctional fiber robots can not only improve current interventional surgeries, such as PCI and AF ablation, but also help provide new therapies to other localized sites. For instance, the magnetically steerable capability enables guidance to the clogged site during PCI and angioplasty in microvessels while the delivered laser can remove the blockage in the artery^56^. Based on their electrophysiological recording, bioimpedance sensing, and pacing functions, these fiber robots can potentially provide diagnosis and treatment during AF ablation^57^. Considering the electrochemotherapy is also a potential application of the fiber robots, by increasing the uptake of high localized doses of chemotherapy drugs to post-chemotherapy resistant tumor sites in combination with electric fields^58^. The localized delivery can reduce many negative side effects in patients with healthy, drug-sensitive cells present in the periphery of hard-to-reach tumor sites, especially those showing poor drug response due to previous treatments.

Moreover, the technology we have developed may open new perspectives in the workup and management of brain diseases, such as deep-seated brain tumors and epileptic seizures. Electroporation-based therapy can induce the blood-brain-barrier (BBB) disruption, thus allowing drug or targeted molecules into the brain. These fiber robots provide an alternative to deliver targeted molecules to brain tumor sites that are difficult to resect and reach with current electrode setups, with the additional benefits of BBB disruption and increased localized drug uptake^59,60^. In addition, recording and stimulation of nerves via an endovascular approach can mitigate the risks associated with open brain surgery by significantly reducing the recovery time and site infections^61^. However, the current endovascular neural recording and stimulation devices are limited to several millimeters^62–64^, thus not suitable for working in the minuscule and tortuous cerebral vasculature. Given the miniaturized size and active steering capability, our fiber robots may access deep brain regions through the endovascular path, and perform localized neural electrophysiological recording and stimulation for treating neurological disorders.

For preclinical studies and clinical translation in the future, several approaches can be applied to improve the performance of the MFFR systems. First, the controlling and integration of MFFRs need to be further investigated to adapt human-size magnetic control systems such as Stereotaxis Niobe and Aeon Phocus that can spatially and temporally manipulate the generated magnetic field^65^. Secondly, as the fibers contain a large number of radiopaque particles that are visible under x-ray, the standard fluoroscopy can be directly applied to visualize the fiber robots in clinic^66^. Ultrasound and photoacoustic imaging can also be alternative options for imaging. Besides, because the MFFRs can also serve as miniature endoscopies, the images transported from the fiber tip can also help indicate the location of the fiber robots. Furthermore, when combined with the lab-on-fiber techniques^67^ and assembled with more advanced and complicated structures, such as semiconductor diodes and sensors^68^, nanoplasmonic structures^69^, and fiber Bragg grating^70^, our MFFRs can enable more functionalities including multimodal biochemical and physiological sensing. Finally, based on the MFFR technology we developed, we can build a system that includes acquisition and analysis programs and even multiple fibers to meet any requirement for different purposes such as simultaneously monitoring in different organs, and closed-loop real-time diagnosis and therapy. Overall, the proposed multifunctional fiber robotics can open avenues to targeted multimodal detection and treatment for hard-to-reach surgical locations in a minimally invasive and remotely controllable manner.

## Methods

### NdFeB composite preparation

NdFeB composite was prepared by thermally mixing SEBS (G1657, Kraton) and NdFeB microparticles with an average diameter of 5 μm (MQFP-B+, Magnequench). SEBS pellets were pressed into sheets at 180 °C using a hot press (MTI Corporation) and the sheets were weighted. NdFeB particles were weighted for desired volume percentage loading and then sprinkled between SEBS sheets. Then, the SEBS sheets with deposited NdFeB were pressed in a hot press at 180 °C and 50 bar for 10 min to embed the NdFeB particles into SEBS sheets. To make a homogeneous dispersion of NdFeB particles in SEBS, the obtained sheets were folded and pressed at the same conditions for 8-10 cycles.

### Multifunctional fiber fabrication

Many fiber structures have been prepared in this study.

For the fiber F1 in Figure 2, a thin layer of PMMA (Rowland Technologies) and a relatively thicker layer of polycarbonate were added to a polycarbonate rod (McMaster-Carr) and consolidated at 180 °C in a vacuum oven. Then two grooves were machined on the thicker polycarbonate layer while one of them was filled by the BiSn electrode. A thin layer of polycarbonate, a thick layer of ferromagnetic composite (FC) layer, and another layer of PC were wrapped around the rod and the whole preform was consolidated again in a vacuum oven.

For the fiber F2 in Figure 2, a thin layer of PMMA (Rowland Technologies) and a relatively thicker layer of polycarbonate were added to a polycarbonate rod (McMaster-Carr) and consolidated at 180 °C in a vacuum oven. Then four grooves were machined on the thicker polycarbonate layer. A thin layer of polycarbonate, a thick layer of ferromagnetic composite (FC) layer, and another layer of polycarbonate were wrapped around the rod and the whole preform was consolidated again in a vacuum oven.

For the fiber F3 in Figure 2, two grooves were machined on a polycarbonate tube. After that, a thin layer of polycarbonate was wrapped around the tube, followed by a FC layer, and a polycarbonate layer and put in a vacuum oven for consolidation.

For the fiber F4 in Figure 2, four grooves were machined on a polycarbonate tube. Then, a thin layer of polycarbonate, a FC layer, and another polycarbonate layer were wrapped on the tubing and then consolidated.

For the fiber used for imaging in Figure 7, 320 PMMA optical fibers with a diameter of 500 μm (SK-20, Industrial Fiber Optics) were filled in a PMMA tube (McMaster-Carr), which was then followed by the wrapping of a FC layer, and a PMMA layer. The preform was then consolidated at 140 °C in a vacuum oven.

All the preforms were drawn into fibers using a custom-built fiber drawing tower. For convergence drawing, two silver wires with a diameter of 100 μm (Surepure Chemetals) and a silica optical fiber (AFH105/125/145T, Fiberguide) were fed through the grooved holes of the preforms. After thermal drawing, the outer polycarbonate or PMMA sacrificial layer was etched using acetone (Sigma-Aldrich) with the assistance of ultrasonic. The Ag/AgCl electrodes were obtained by dipping fiber with Ag electrodes into 0.1-M FeCl3 solution for 1 min.

### Magnetic and mechanical characterization

The magnetization of the NdFeB composite as a function of magnetic field was measured using a vibrating sample magnetometer (MicroSense EZ9), equipped with an electromagnet with a maximum output field of 2.2 T. Samples of composites before and after the thermal drawing process were measured. The samples before the drawing process were 0.5 mm thick square sheets with lateral areas of approximately 40 mm^2^; the magnetic field was applied parallel to the square’s edge. The samples after the drawing process were cylindrical fibers cut to approximately 8 mm long segments; the magnetic field was applied along the axis of the fiber. For each magnetometry measurement, the sample was attached to a Pyrex rod. The diamagnetic background was subtracted, and the measured magnetization was normalized by the volume of the composite. The electromagnet of the magnetometer was also used to magnetize the fibers used in this study.

Mechanical stress-strain tests were measured using a dynamic mechanical analysis (DMA Q800, TA instruments). The heat-pressed NdFeB particles loaded SEBS sheets were cut into rectangular stripes. The Young’s moduli were identified by fitting the experimental curve into a neo-Hookean model.

### Magnetic actuation and in vitro demonstration

Magnetic actuation characterization was performed under a pair of custom-made Helmholtz coils. The coils were produced by winding polyamide-coated copper magnet wire (Remington Industries) onto polycarbonate tubes (d = 7.5cm, McMaster-Carr). The coils were powered by a DC power supply (current 0-14A, TDK Lambda). The generated uniform magnetic field between the two coils was in the range of 0-45 mT. The relationship between the applied current and generated uniform magnetic field can be described as *B* = *I* * 3.63 *mT*/*A*. The fiber samples were fixed on a holder and placed perpendicular to the magnetic field. In order to reduce the influence of temperature increase due to Joule heating, ice bags were used for cooling between the measurements. Each time before the measurement, the uniform magnetic field density was verified using a commercial magnetic field meter (TM-197, Temmars). The displacement of the fiber samples was recorded using a camera.

For all demonstrations presented in the paper, a permanent magnet (diameter and height of 50 mm; DY0Y0-N52, K&J Magnetics Inc.) was used to apply the desired magnetic fields for actuation at a certain distance. The desired strength and direction of the applied magnetic field were controlled by manually manipulating the position and orientation of the magnet.

To reduce the friction between fiber and phantom, we can coat a hydrogel skin to the fibers following the protocol reported previously (*20*). The cleaned samples were immersed in an acetone solution with 10 wt% benzophenone (Sigma-Aldrich) for 15 minutes. Both ends of the fibers were sealed with 5-min epoxy (Loctite) to protect the polycarbonate and PMMA layer. Then the samples were immersed in a pre-gel solution containing 30 wt % hydrogel monomers (N,N′-dimethylacrylamide; DMAA, Sigma-Aldrich) and 1 wt % Irgacure 2959 (Sigma-Aldrich) based on deionized water and the solution was subjected to UV irradiation for 60 minutes. The unreacted regents were removed by rinsing the sample with a large amount of deionized water for 48 hours.

The 2D vessel phantoms used for demonstration in Fig. 3A, 3B was modeled with computer-aided design software (Solidworks) and manufactured by machining a polycarbonate plate using CNC. A polycarbonate cover (thickness: 2 mm) was placed on the top of the phantom to seal it. The 3D silicone vessel phantom in Fig. 3C was purchased from Trandomed. During the demonstration, the phantoms were filled with 22 % (v/v) glycerol as a blood mimic. The human heart model in Fig. 4A was purchased from Axis Scientific. The brain phantom was fabricated using 0.6 wt% agarose gel.

### Electroporation testing

For electroporation testing, we manufactured a 4-well mold consisting of 10 mm diameter wells on a PLA 3D printer. PDMS was used to attach glass coverslips to the bottom of the wells to enable easy imaging of cells and fibers in the gel. We incorporated fibers into the molds from the sidewall, fastened with PDMS to prevent any movement from potential lesion site when imaging.

We applied 100μs monopolar pulses to 3D in-vitro hydrogel platforms to demonstrate the fiber platform’s ability to electroporate cells. We measured lesion areas 30 minutes after pulsing from fluorescent microscopy images of ablated regions adjacent to the electrodes using ZEN Blue (Carl Zeiss®). We demonstrate the effect of conventional IRE treatments in 5mg/ml gels combined with a calcium adjuvant on U-251 tumor cells. The electroporated areas are defined by YO-PRO-1 uptake (green) and live cells with no membrane disruption are marked by Calcein Red-AM. We applied 150 V to produce fields capable of electroporation at the fiber’s surface, limited by the onset of arcing which would destroy the fibers.

We cultured U-251 glioblastoma cells (Sigma-Aldrich, St. Louis, MO) in Dulbecco’s modified eagle medium (DMEM, ATCC) supplemented with 10% fetal bovine serum (FBS, ATCC) and 1% penicillin-streptomycin (Fisher Scientific, Suwanee, GA). Cells were cultured in 75 cm^2^ flasks and were maintained in an incubator with 5% CO2 and 100% humidity at 37°C. Cells were monitored and sub-cultured regularly during the exponential phase to maintain viability and to avoid overgrowing. We incorporated these cells into hydrogel constructs upon reaching confluency. We dissolved lyophilized collagen in sterile 0.1% acetic acid solution in deionized water to obtain a concentration of 10 mg/ml. Collagen provides a convenient scaffold material that produces relevant 3D geometry, integrin engagement with the surrounding extracellular matrix, and appropriate cell-cell interactions. The dissolved collagen was mixed with 10x DMEM and 1N NaOH to attain a pH of 7.4 and a final density of 5 mg/mL. We then introduced U-251 cells into the collagen mixture, homogenized them, and injected them into prefabricated, sterile PDMS wells. We incubated the molds after gel addition at 37°C for 25 minutes to allow the collagen to fully polymerize. 1 mL of media was added to each well and the mold was placed in a covered petri dish and incubated at 37 °C for a minimum of 24 hours before testing.

We utilized a BTX 830 ECM Electroporator (Harvard Apparatus ©) to electroporate our tumor mimics with 200 rectangular pulses of 100 μs length at voltages of 175 V at a frequency of 1 Hz delivered through the fiber. We captured the waveforms delivered using a high voltage probe (Enhancer 300, BTX, Holliston, MA) connected to an oscilloscope (LeCroy Waverunner 4zi) to capture the waveform applied. For Reversible tests, we exposed the gels to fluorescent dyes for 30 minutes before treatment. For IRE tests, staining was done 24 hours post-treatment. We added 2mM of Calcein Red-AM (ThermoFisher Scientific) and 1μM YO-PRO-1 (ThermoFisher Scientific) to 400 uL of PBS in each well. Wells were triple washed with PBS before imaging for IRE tests (24-hour time point) and we aspirated this staining solution before RE treatment. We used a 1 mL syringe with a needle to introduce 5mM Ca^2+^ through the drug delivery port between the electrodes. Immediately upon pumping the drug, we delivered the treatment to the gels.

Each well was imaged immediately after the treatment. Gels were rinsed with Dulbecco’s phosphate-buffered saline prior to being imaged on a confocal microscope (Carl Zeiss, at 100x total magnification (10x objective, 10x eyepiece)). We stitched tile scans on the Zen Blue software (Carl Zeiss) to encompass the region around the fiber’s surface. 20 stacks were taken to cover the top and bottom of the fiber. Gel images were projected onto a 2D area to measure the lateral lesion areas. The Green (YO-PRO-1 uptake) signal was the primary measure of the electroporated region. We projected the z-stacks to obtain the maximum 2D area distribution and correlated these areas with EF isolines obtained from COMSOL using a curve fitting software (MATLAB, Natick, MA).

Due to a combination of small lesion volume, minimal cells present within the electroporated area (despite increasing cell density) and low resolution of z movement offered by the confocal microscope, the electroporated regions are small relative to cell size to ensure greater accuracy of measurement.

### Numerical Model

We modeled our fiber as a 200 μm long fiber with a diameter of 500 μm in a gel construct 1 mm in diameter and 500 μm thick in COMSOL Multiphysics, (v6.0). We built a model of the fiber probe, with 2 silver wires enclosed by a polycarbonate cylinder which has a drug delivery port in the center placed in this hydrogel scaffold. With the walls insulated, the effective electrode surface exposed to the gel behaves like 2 cylindrical discs 100 μm in diameter(each). The model determines the electric field distribution in the hydrogel was solved by the Laplace equation while accounting for the Joule heating during the treatment to estimate the temperature change (and its impact on conductivity and field distribution). The temperature was calculated by the heat transfer equation as previous reported^71^. We measured the electric field threshold of the electroporation by correlating the projected maximum area of the lesion with the isolines obtained for the applied electric field.

### Fiber bundle imaging

The masks were prepared through laser marking. We used spray paint to form a thin layer of paint on a glass slide. Then patterns were directly written by removing paint via a programmed laser beam using a laser marker system (TM-station, Boss Laser). For the imaging setup, a halogen lamp (Ocean Optics) was used as the broadband light source. Then the illuminated light was focused on the custom-made mask and the image of the pattern on the mask was then formed on the fiber end through a lens set. The focal lengths of the two lenses were 40 mm and 48 mm, respectively, resulting in a magnification of 1.2. The imaging from the other end of the fiber was collected by a CCD camera (Stingray F145B ASG, Allied) through an objective. Both ends of the imaging fiber were polished for better coupling efficiency.

### Electrochemical impedance spectrum for neural probes

The fiber probes (fiber F1) of two centimeters were prepared and the inner electrode was electrically connected to the copper wire. The electrochemical impedance measurement was performed via a potentiostat (Interface 1010E, Gamry Instruments). During the measurements, the fiber probes were inserted in 1x phosphate-buffered saline (PBS, Thermo Fisher) as working electrodes, while a Pt electrode served as a reference and counter electrode with applied AC voltage sweep (10mV, 100-10kHz).

### Multifunctional probes assembly

For all fibers used in this project, the TPE jackets at the back end were peeled off for connection. The fiber ends were first inserted into fiber optical ferrules and sealed using phenyl salicylate. To obtain a flat surface for light coupling, the ferrule top parts were polished by optical polishing papers with roughness from 30um to 1um. Then, the electrodes embedded inside the fibers were exposed via a razor blade for electrical connection. Silver paint (SPI Supplies) was applied to the exposed sites individually, followed by wrapped copper wires and another layer of silver paint. The connected copper wires were soldered to the pin connectors (Sullins Connector Solutions) for further connection to electrical stimulators or signal recording setup. Similarly, the microfluidic channels inside the fibers were exposed manually via a razor blade and the fibers were inserted into ethylene-vinyl acetate tubing (0.5 mm inner diameter) with the assistance of a needle. The fibers were placed perpendicular to the tubing and the exposed sites were inside channels of the tubing. Epoxy (LOCTITE) was applied to the edge between fiber and tubing to prevent leakage. Finally, another layer of epoxy was applied to all the electrical and optical connection parts for further fixation.

### Animal experiments

All animal procedures were approved by Virginia Tech Institutional Animal Care and Use Committee and Institutional Biosafety Committee and were carried out in accordance with the National Institutes of Health Guide for the Care and Use of Laboratory Animals.

For the experiments using Langendorff mouse heart models, 25-week C57BL/6J mice were anesthetized with isoflurane inhalation. Hearts were rapidly excised, cannulated, and retrogradely perfused with a crystalloid perfusion solution (in mM: NaCl 139.5, MgCl 1, NaH2PO4 1.2, KCl 4, CaCl2 1.8, Glucose 5.6 and HEPES 10; pH=7.4 adjusted with 5.5-mM NaOH) and then placed in a 3D printed PLA bath filled by the same solution maintaining heart temperature at 36°C. A fiber with a diameter of 600 μm was inserted from the superior vena cava into the right ventricle through the tricuspid valve. The fiber tip was placed against the ventricular wall to make sure that the two Ag/AgCl electrodes of the fiber contacted the cardiac muscle. All the electrical recording or stimulation was performed after a stabilization period of 20-30 minutes. A volume conducted bath ECG was recorded through an ECG amplifier (Hugo Sachs Elektronik) at 1 kHz, while 2 electrode wires were placed on either side of the ventricles and the ground was placed at the rear of the bath. The bipolar EGM was recorded through the two Ag/AgCl electrodes embedded in the fiber via a custom-made differential amplifier circuit. Both the ECG and EGM signals were recorded by a data acquisition system (DAQ, Powerlab) and read using LabChart. During pacing, the stimulation pulses (0.08mA) were delivered from a stimulus isolator (WPI) to the ventricle muscles through the two electrodes inside the fiber. For bioimpedance measurement, two-electrode experiments were performed while one Ag/AgCl electrode inside the fiber served as a working electrode and the other one as a counter and reference electrode by an AC current of 0.03 mA (1 Hz – 100kHz). 26.1g/L mannitol solution was added through the perfusion system. Bioimpedance measurements were performed right before the mannitol perfusion and 20 minutes after the perfusion.

For the mouse brain-related experiments, 10-12 week old C57BL/6J (JAX#000664) x FVBN/J (JAX#001800) mice were bred from C57BL/6J (JAX#000664) and FVBN/J (JAX#001800) received from the Jackson Laboratory (Bar Harbor, ME). Mice had access to food and water ad libitum and were kept in a facility maintained for a 12-h light/dark cycle starting at 7 AM. Mice were set up on a stereotaxic apparatus (RWD) and 1–3.5% isoflurane was induced to animals via nose cone during all procedures for anesthesia. To expose the skull, an incision was made on the skin along the midline and then all tissue was removed from the surface of the skull. Optibond (Kerr Dental) was applied to the skull to prime it for dental cement. A craniotomy was made above the cerebellum, and a stainless-steel ground wire was implanted just below the skull. Additionally, a titanium head-bar was implanted above the cerebellum. Mice were habituated to head fixation over the course of one week. After habituation, a craniotomy was made above dorsal CA1 (coordinate, in mm from bregma: −1.75 posterior, 1.5 mm left). While head-fixed above a running wheel, the fiber probe was lowered in CA1. Neural data were acquired at 30 kHz using an Intan RHD2000 recording system.

## Supplementary Materials

Supplementary text

Figs. S1 to S7

Movies S1 to S4

## Author contributions

Y.Z. and X.J. conceived and designed the study. Y.Z. developed the ferromagnetic fiber robots. Y.Z., Y.Lim, Y.Li and R.W. characterized the fiber robots, X.W. and Y.Z. conducted the Langendorff-perfused heart experiments and analyzed the results. R.A.V., Y.Z. and K.D. conducted the in vitro electroporation experiments and analyzed the results, J.K., Y.Z., E.G. and S.J. conducted the in vivo animal study and analyzed the results. Y.Z. and X.J. wrote the manuscript with input from all authors. X.J., S.P., R.V.D., S.E., D.E., H.S. and A.W. supervised the study.

## Data and materials availability

All data are available in the main text or the supplementary materials.

## References and Notes

1 Li, M., Pal, A., Aghakhani, A., Pena-Francesch, A. & Sitti, M. Soft actuators for real-world applications. Nature Reviews Materials, 1–15 (2021).

2 Garcia, L. et al. The Role of Soft Robotic Micromachines in the Future of Medical Devices and Personalized Medicine. Micromachines 13, 28 (2022).

3 Cianchetti, M., Laschi, C., Menciassi, A. & Dario, P. Biomedical applications of soft robotics. Nature Reviews Materials 3, 143–153 (2018).

4 Yang, G.-Z. et al. The grand challenges of Science Robotics. Science robotics 3, eaar7650 (2018).

5 Runciman, M., Darzi, A. & Mylonas, G. P. Soft robotics in minimally invasive surgery. Soft robotics 6, 423–443 (2019).

6 Rafii-Tari, H., Payne, C. J. & Yang, G.-Z. Current and emerging robot-assisted endovascular catheterization technologies: a review. Annals of biomedical engineering 42, 697–715 (2014).

7 Piskarev, Y. et al. A Variable Stiffness Magnetic Catheter Made of a Conductive Phase-Change Polymer for Minimally Invasive Surgery. Advanced Functional Materials, 2107662 (2022).

8 Menaker, S. A. et al. Current applications and future perspectives of robotics in cerebrovascular and endovascular neurosurgery. Journal of neurointerventional surgery 10, 78–82 (2018).

9 Li, J., Ávila, B. E.-F. d., Gao, W., Zhang, L. & Wang, J. Micro/nanorobots for biomedicine: Delivery, surgery, sensing, and detoxification. Science Robotics 2, eaam6431 (2017). https://doi.org:doi:10.1126/scirobotics.aam6431

10 Weisz, G. et al. Safety and feasibility of robotic percutaneous coronary intervention: PRECISE (Percutaneous Robotically-Enhanced Coronary Intervention) Study. Journal of the American College of Cardiology 61, 1596–1600 (2013).

11 Han, M. et al. Submillimeter-scale multimaterial terrestrial robots. Science Robotics 7, eabn0602 (2022). https://doi.org:doi:10.1126/scirobotics.abn0602

12 Wu, Z. et al. A microrobotic system guided by photoacoustic computed tomography for targeted navigation in intestines in vivo. Science Robotics 4, eaax0613 (2019). https://doi.org:doi:10.1126/scirobotics.aax0613

13 Overvelde, J. T., Kloek, T., D’haen, J. J. & Bertoldi, K. Amplifying the response of soft actuators by harnessing snap-through instabilities. Proceedings of the National Academy of Sciences 112, 10863–10868 (2015).

14 Kim, Y., Yuk, H., Zhao, R., Chester, S. A. & Zhao, X. Printing ferromagnetic domains for untethered fast-transforming soft materials. Nature 558, 274–279 (2018).

15 Hu, W., Lum, G. Z., Mastrangeli, M. & Sitti, M. Small-scale soft-bodied robot with multimodal locomotion. Nature 554, 81–85 (2018).

16 Ze, Q. et al. Magnetic shape memory polymers with integrated multifunctional shape manipulation. Advanced Materials 32, 1906657 (2020).

17 Kim, Y. et al. Telerobotic neurovascular interventions with magnetic manipulation. Science Robotics 7, eabg9907 (2022).

18 Hwang, J. et al. An Electromagnetically Controllable Microrobotic Interventional System for Targeted, Real - Time Cardiovascular Intervention. Advanced Healthcare Materials, 2102529 (2022).

19 Zhao, R., Kim, Y., Chester, S. A., Sharma, P. & Zhao, X. Mechanics of hard-magnetic soft materials. Journal of the Mechanics and Physics of Solids 124, 244–263 (2019).

20 Wang, L., Kim, Y., Guo, C. F. & Zhao, X. Hard-magnetic elastica. Journal of the Mechanics and Physics of Solids 142, 104045 (2020).

21 Kim, Y., Parada, G. A., Liu, S. & Zhao, X. Ferromagnetic soft continuum robots. Science Robotics 4, eaax7329 (2019).

22 Cui, J. et al. Nanomagnetic encoding of shape-morphing micromachines. Nature 575, 164–168 (2019).

23 Deng, H. et al. Laser reprogramming magnetic anisotropy in soft composites for reconfigurable 3D shaping. Nature communications 11, 1–10 (2020).

24 Alapan, Y., Karacakol, A. C., Guzelhan, S. N., Isik, I. & Sitti, M. Reprogrammable shape morphing of magnetic soft machines. Science advances 6, eabc6414 (2020).

25 Pancaldi, L. et al. Flow driven robotic navigation of microengineered endovascular probes. Nature communications 11, 1–14 (2020).

26 Zhou, C. et al. Ferromagnetic soft catheter robots for minimally invasive bioprinting. Nature communications 12, 1–12 (2021).

27 Loke, G., Yan, W., Khudiyev, T., Noel, G. & Fink, Y. Recent progress and perspectives of thermally drawn multimaterial fiber electronics. Advanced Materials 32, 1904911 (2020).

28 Guo, Y. et al. Miniature multiplexed label-free pH probe in vivo. Biosensors and Bioelectronics 174, 112870 (2021).

29 Zhang, Y. et al. Thermally drawn stretchable electrical and optical fiber sensors for multimodal extreme deformation sensing. Advanced Optical Materials 9, 2001815 (2021).

30 Canales, A. et al. Multifunctional fibers for simultaneous optical, electrical and chemical interrogation of neural circuits in vivo. Nature biotechnology 33, 277–284 (2015).

31 Koppes, R. A. et al. Thermally drawn fibers as nerve guidance scaffolds. Biomaterials 81, 27–35 (2016).

32 Jin, Y. et al. Functional skeletal muscle regeneration with thermally drawn porous fibers and reprogrammed muscle progenitors for volumetric muscle injury. Advanced Materials 33, 2007946 (2021).

33 Park, S. et al. One-step optogenetics with multifunctional flexible polymer fibers. Nature neuroscience 20, 612–619 (2017).

34 Jiang, S. et al. Spatially expandable fiber-based probes as a multifunctional deep brain interface. Nature communications 11, 1–14 (2020).

35 Park, J. et al. In situ electrochemical generation of nitric oxide for neuronal modulation. Nature nanotechnology 15, 690–697 (2020).

36 Qu, Y. et al. Superelastic multimaterial electronic and photonic fibers and devices via thermal drawing. Advanced Materials 30, 1707251 (2018).

37 Yun, S. H. & Kwok, S. J. Light in diagnosis, therapy and surgery. Nature biomedical engineering 1, 1–16 (2017).

38 Bakker, J. M. Electrogram recording and analyzing techniques to optimize selection of target sites for ablation of cardiac arrhythmias. Pacing and Clinical Electrophysiology 42, 1503–1516 (2019). https://doi.org:10.1111/pace.13817

39 Sperelakis, N. & Hoshiko, T. Electrical Impedance of Cardiac Muscle. Circulation Research 9, 1280–1283 (1961). https://doi.org:10.1161/01.res.9.6.1280

40 Cinca, J. et al. Changes in myocardial electrical impedance induced by coronary artery occlusion in pigs with and without preconditioning: correlation with local ST-segment potential and ventricular arrhythmias. Circulation 96, 3079–3086 (1997).

41 Dzwonczyk, R. et al. Myocardial Electrical Impedance Responds to Ischemia and Reperfusion in Humans. IEEE Transactions on Biomedical Engineering 51, 2206–2209 (2004). https://doi.org:10.1109/tbme.2004.834297

42 Veeraraghavan, R., Salama, M. E. & Poelzing, S. Interstitial volume modulates the conduction velocity-gap junction relationship. American Journal of Physiology-Heart and Circulatory Physiology 302, H278–H286 (2012).

43 Neshatvar, N., Regnacq, L., Jiang, D., Wu, Y. & Demosthenous, A. in 2020 IEEE International Symposium on Circuits and Systems (ISCAS). 1–4 (IEEE).

44 Feltes, T. F. et al. Indications for cardiac catheterization and intervention in pediatric cardiac disease: a scientific statement from the American Heart Association. Circulation 123, 2607–2652 (2011).

45 Crawford, K., LeBerge, M., Simionescu, D. & Nagarajan, N. Pediatric Cardiac Devices: Recent Progress and Remaining Problems. (2022).

46 Lang, D., Sulkin, M., Lou, Q. & Efimov, I. R. Optical mapping of action potentials and calcium transients in the mouse heart. JoVE (Journal of Visualized Experiments), e3275 (2011).

47 Martin, C. A., Guzadhur, L., Grace, A. A., Lei, M. & Huang, C. L.-H. Mapping of reentrant spontaneous polymorphic ventricular tachycardia in a Scn5a+/− mouse model. American Journal of Physiology-Heart and Circulatory Physiology 300, H1853–H1862 (2011).

48 George, S. A. et al. Extracellular sodium dependence of the conduction velocity-calcium relationship: evidence of ephaptic self-attenuation. American Journal of Physiology-Heart and Circulatory Physiology 310, H1129–H1139 (2016).

49 Ivey, J. et al. Characterization of ablation thresholds for 3D-cultured patient-derived glioma stem cells in response to high-frequency irreversible electroporation. Research 2019(2019).

50 Wasson, E. M. et al. Understanding the role of calcium-mediated cell death in high-frequency irreversible electroporation. Bioelectrochemistry 131, 107369 (2020).

51 Frank, J. A., Antonini, M.-J. & Anikeeva, P. Next-generation interfaces for studying neural function. Nature biotechnology 37, 1013–1023 (2019).

52 Kozai, T. D. Y. et al. Reduction of neurovascular damage resulting from microelectrode insertion into the cerebral cortex using in vivo two-photon mapping. Journal of neural engineering 7, 046011 (2010).

53 Hong, A., Petruska, A. J., Zemmar, A. & Nelson, B. J. Magnetic control of a flexible needle in neurosurgery. IEEE Transactions on Biomedical Engineering 68, 616–627 (2020).

54 Petruska, A. J. et al. in 2016 IEEE International Conference on Robotics and Automation (ICRA). 4392–4397 (IEEE).

55 Buzsáki, G. et al. Hippocampal network patterns of activity in the mouse. Neuroscience 116, 201–211 (2003). https://doi.org:10.1016/s0306-4522(02)00669-3

56 Dahm, J. B. et al. Laser - facilitated thrombectomy: a new therapeutic option for treatment of thrombus-laden coronary lesions. Catheterization and cardiovascular interventions 56, 365–372 (2002).

57 Wood, M. A., Brown-Mahoney, C., Kay, G. N. & Ellenbogen, K. A. Clinical outcomes after ablation and pacing therapy for atrial fibrillation: a meta-analysis. Circulation 101, 1138–1144 (2000).

58 Wright, J. A. et al. DNA amplification is rare in normal human cells. Proceedings of the National Academy of Sciences 87, 1791–1795 (1990).

59 Sharma, P. et al. Multireceptor targeting of glioblastoma. Neuro-oncology advances 2, vdaa107 (2020).

60 Lorenzo, M. F. et al. Temporal characterization of blood–brain barrier disruption with high-frequency electroporation. Cancers 11, 1850 (2019).

61 Fan, J. Z., Lopez-Rivera, V. & Sheth, S. A. Over the horizon: the present and future of endovascular neural recording and stimulation. Frontiers in Neuroscience 14, 432 (2020).

62 Chen, J. C. et al. A wireless millimetric magnetoelectric implant for the endovascular stimulation of peripheral nerves. Nature Biomedical Engineering, 1–11 (2022).

63 Oxley, T. J. et al. Minimally invasive endovascular stent-electrode array for high-fidelity, chronic recordings of cortical neural activity. Nature biotechnology 34, 320–327 (2016).

64 Opie, N. L. et al. Focal stimulation of the sheep motor cortex with a chronically implanted minimally invasive electrode array mounted on an endovascular stent. Nature biomedical engineering 2, 907–914 (2018).

65 Hwang, J., Kim, J.-y. & Choi, H. A review of magnetic actuation systems and magnetically actuated guidewire-and catheter-based microrobots for vascular interventions. Intelligent Service Robotics 13, 1–14 (2020).

66 Pané, S. et al. Imaging technologies for biomedical micro - and nanoswimmers. Advanced Materials Technologies 4, 1800575 (2019).

67 Vaiano, P. et al. Lab on Fiber Technology for biological sensing applications. Laser & Photonics Reviews 10, 922–961 (2016).

68 Rein, M. et al. Diode fibres for fabric-based optical communications. Nature 560, 214–218 (2018).

69 Jiang, S. et al. Nano-optoelectrodes integrated with flexible multifunctional fiber probes by high-throughput scalable fabrication. ACS Applied Materials & Interfaces 13, 9156–9165 (2021).

70 Cusano, A., Cutolo, A. & Albert, J. Fiber Bragg grating sensors: recent advancements, industrial applications and market exploitation. (Bentham Science Publishers, 2011).

71 Kim, J. et al. Laser Machined Fiber-Based Microprobe: Application in Microscale Electroporation. Advanced Fiber Materials (2022). https://doi.org:10.1007/s42765-022-00148-5

