## supplementary material for "Multifunctional ferromagnetic fiber robots for navigation, sensing, and treatment in minimally invasive surgery"

**This PDF file includes:**

Supplementary Text: Supplementary notes 1-2

Figs. S1 to S6

Captions for Movies S1 to S4

**Other Supplementary Materials for this manuscript include the following:**

Movies S1 to S4

### List of contents:

Supplementary Note 1. Characterization of NdFeB doped SEBS.

Supplementary Note 2. Analytical model for ferromagnetic fiber robots.

Fig. S1 Illustration of preform preparation process for fiber F1. A thin layer of PMMA and a relatively thicker layer of PC are added to a PC rod and consolidated at 180 °C in a vacuum oven. Then two grooves are machined on the thicker PC layer while one of them is filled by the BiSn electrode. A thin layer of PC, a thick layer of NdFeB particles doped SEBS and another layer of PC are wrapped around the rod and the whole preform is consolidated again in a vacuum oven.

Fig. S2 Characterization of NdFeB doped SEBS. (A) Young's modulus (denoted $E_{eff}$ ) of the ferromagnetic composite at different particle fractions. (B) Normalized deflection δ/L predicted from theory plotted against ferromagnetic particle loading volume fraction with different core-to-fiber ratios: d/D = 0.42, 0.52 0.57, 0.64 when L/D = 40. (C) Particle loading volume fraction when the fiber defections reach the highest value predicted from theory against core-to-fiber ratios.

Fig. S3 Photography of a spool with ~150 m long continuous ferromagnetic fiber (F1) drawn from one preform.

Fig. S4 Sideview image of the commercial guidewire (size, 0.36 mm) and fiber robot (0.38 mm) used for in vivo evalulation in 2D vascular phantom (Fig. 3A,B).

Fig. S5 Energy-dispersive X-ray spectroscopy for the fiber surface shown in Fig. 3d(ii). A 10nm Pt layer is coated on the fiber surface using sputter for increasing the conductivity of the fiber and getting better SEM imaging. Ag and Cl elements show highest peak from the spectrum. The C and O elements can be from polycarbonate particles generated from surrounding polymer layers during SEM sample preparation process. The Fe elements can be from $FeCl_3$ residuals from $FeCl_3$ solution.

Fig. S6 Impedance spectra of perfusion solution measured through the fiber before and after mannitol perfusion. The solution impedance stays consistent after mannitol perfusion.

Fig. S7 Characterization of optical properties of neural probe fiber (fiber F1) and image guide fiber. (A) The optical transmission spectrum obtained at a wavelength range of 450 – 750 nm for neural probe fiber (normalized to the maximum value). (B) The transmission loss at a wavelength of 473 nm measured using cut back method is 0.797 dB/cm ($R^2$=0.99592) for neural probe fiber. (C) The optical transmission spectrum obtained at a wavelength range of 450 – 750 nm for image guide fiber (normalized to the maximum value). (D) The transmission loss at a wavelength of 615 nm measured using cut back method is 0.316 dB/cm ($R^2$=0.9915) for image guide fiber.

Movie S1 In vitro evaluation of commercial guidewire in 2D vascular phantom.

Movie S2 In vitro evaluation of MFFR in 2D vascular phantom.

Movie S3 In vitro evaluation of MFFR in 3D vascular phantom.

Movie S4 Demonstration of MFFR in an artificial human heart model.

**Supplementary Note 1. Characterization of NdFeB doped SEBS.**

The Young's modulus of the doped SEBS increases nonlinearly as the NdFeB particle increases (**Supplementary Fig. 2A**), which can be described using the Mooney Equation with the assumption that the particles are close packed spheres, and the composites are incompressible[1],

$$E_{jacket} = E_0 \exp\left(\frac{2.5\phi}{1 - 1.35\phi}\right) \tag{S1}$$

where $E_0$denote the Young's modulus of undoped SEBS, which is 2.1 MPa.

**Supplementary Note 2. Analytical model for ferromagnetic fiber robots.**

Based on the method developed by Wang, et.al[2], the governing equation can be simplified as

$$\frac{E_{eff}I}{A}\frac{d^2\theta}{ds^2} + M_{eff}Bcos\theta = 0 \tag{S2}$$

The angular displacement at the free end of the fiber $\theta_L$ can be obtained by solving

$$L = \sqrt{\frac{E_{eff}I}{2M_{eff}BA}}\int_0^{\theta_L}\frac{d\theta}{\sqrt{sin\theta_L - sin\theta}} \tag{S3}$$

The deflection of the free end of the fiber along the y-axis is

$$\delta = \sqrt{\frac{E_{eff}I}{2M_{eff}BA}}\int_0^{\theta_L}\frac{sin\theta d\theta}{\sqrt{sin\theta_L - sin\theta}} \tag{S4}$$


1 Ahmed, S. & Jones, F. A review of particulate reinforcement theories for polymer composites. *Journal of materials science* **25**, 4933-4942 (1990).

2 Wang, L., Kim, Y., Guo, C. F. & Zhao, X. Hard-magnetic elastica. *Journal of the Mechanics and Physics of Solids* **142**, 104045 (2020).

### Supplementary Figures

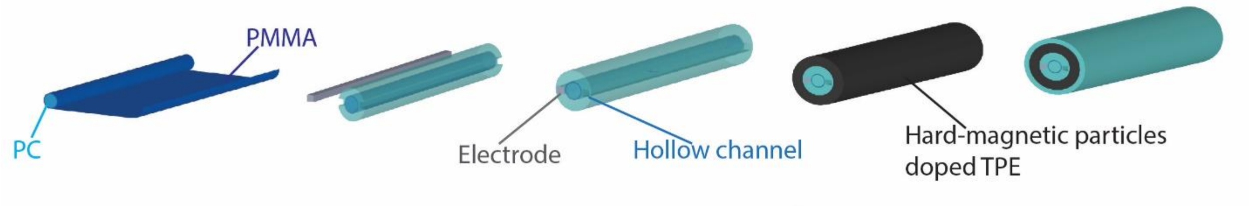


**Fig. S1 Illustration of preform preparation process for fiber F1.** A thin layer of PMMA and a relatively thicker layer of PC are added to a PC rod and consolidated at 180 ºC in a vacuum oven. Then two grooves are machined on the thicker PC layer while one of them is filled by the BiSn electrode. A thin layer of PC, a thick layer of NdFeB particles doped SEBS, and another layer of PC are wrapped around the rod and the whole preform is consolidated again in a vacuum oven.

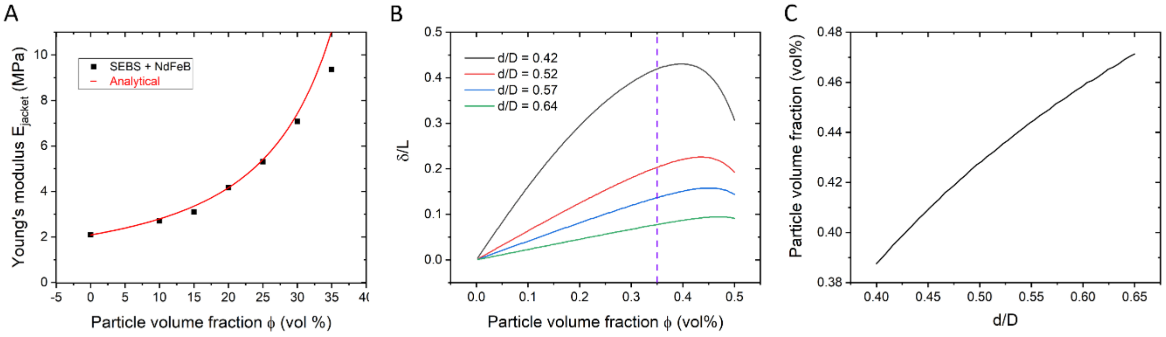


**Fig. S2 Characterization of NdFeB doped SEBS.** (**A**) Young's modulus (denoted $E_{eff}$ ) of the ferromagnetic composite at different particle fractions. (**B**) Normalized deflection δ/L predicted from theory plotted against ferromagnetic particle loading volume fraction with different core-to-fiber ratios: d/D = 0.42, 0.52 0.57, 0.64 when L/D = 40. The fiber deflections increase with the loading fraction when the fraction is below 35 volume%. (**C**) Particle loading volume fraction when the fiber defections reach the highest value predicted from theory against core-to-fiber ratios.

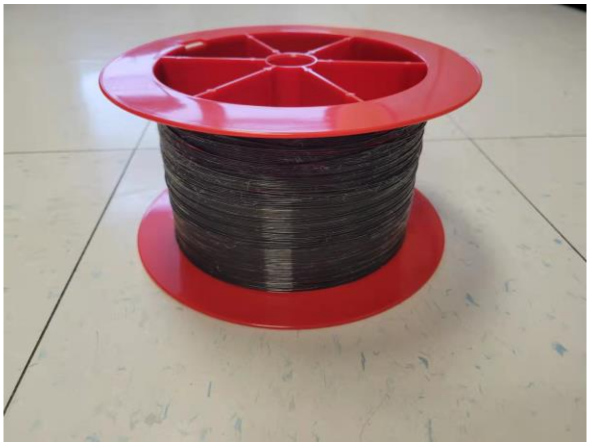

**Fig. S3 Photography of a spool with ~150m long continuous ferromagnetic fiber (F1) drawn from one preform.** The fiber has one waveguide, one BiSn electrode, and one hollow channel. The size of the fiber is ~500 µm with a polycarbonate sacrificial layer.

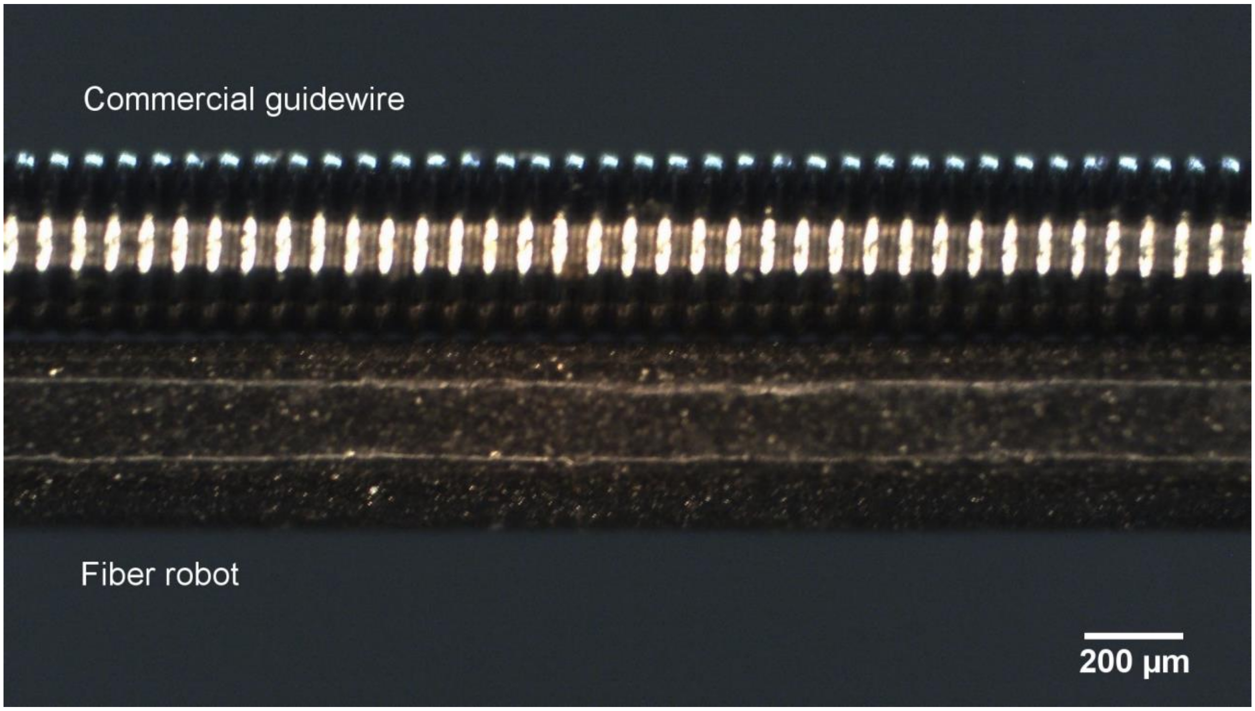


**Fig. S4 Sideview image of the commercial guidewire (size, 0.36 mm) and fiber robot (0.38 mm) used for in vivo evalulation in 2D vascular phantom (Fig. 3A,B).**

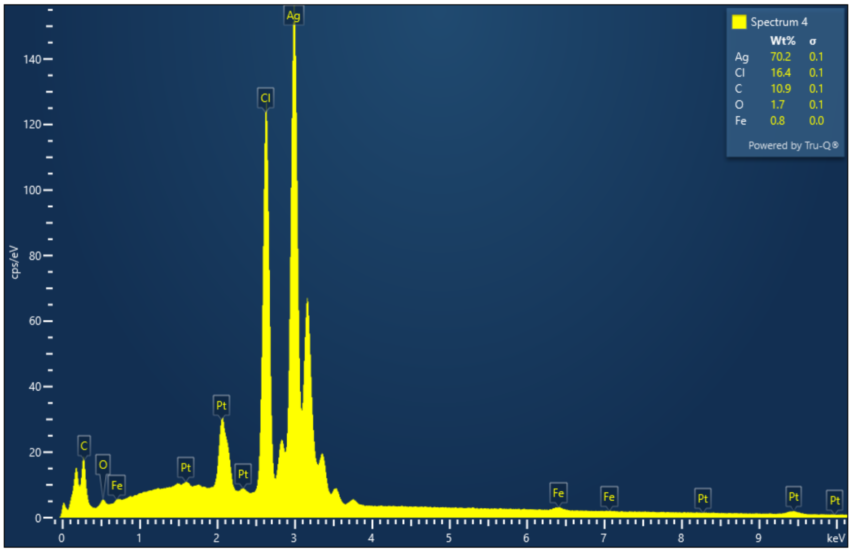


**Fig. S5 Energy-dispersive X-ray spectroscopy for the fiber surface in Fig. 3d(ii).** A 10 nm Pt layer is coated on the fiber surface using sputter for increasing the conductivity of the fiber and getting better SEM imaging. Ag and Cl elements show the highest peak from the spectrum. The C and O elements can be from polycarbonate particles generated from surrounding polymer layers during the SEM sample preparation process. The Fe elements can be from $FeCl_3$ residuals from $FeCl_3$ solution.

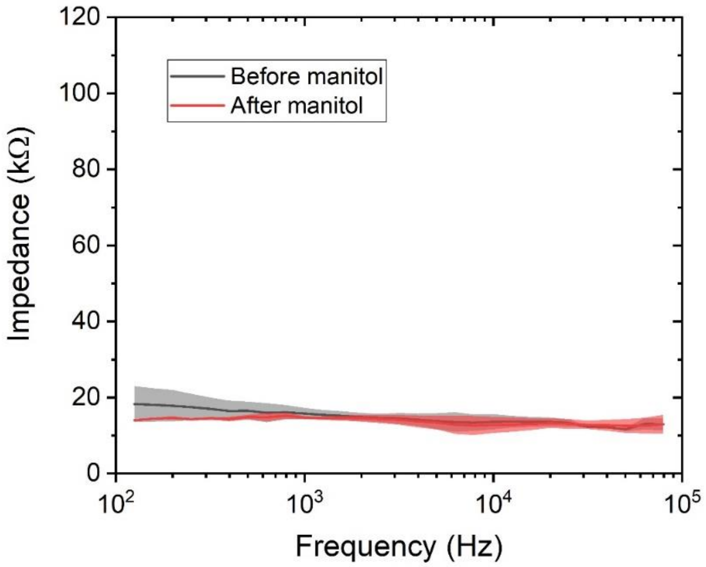


**Fig. S6 Impedance spectra of perfusion solution measured through the fiber before and after mannitol perfusion.** The solution impedance stays consistent after mannitol perfusion.

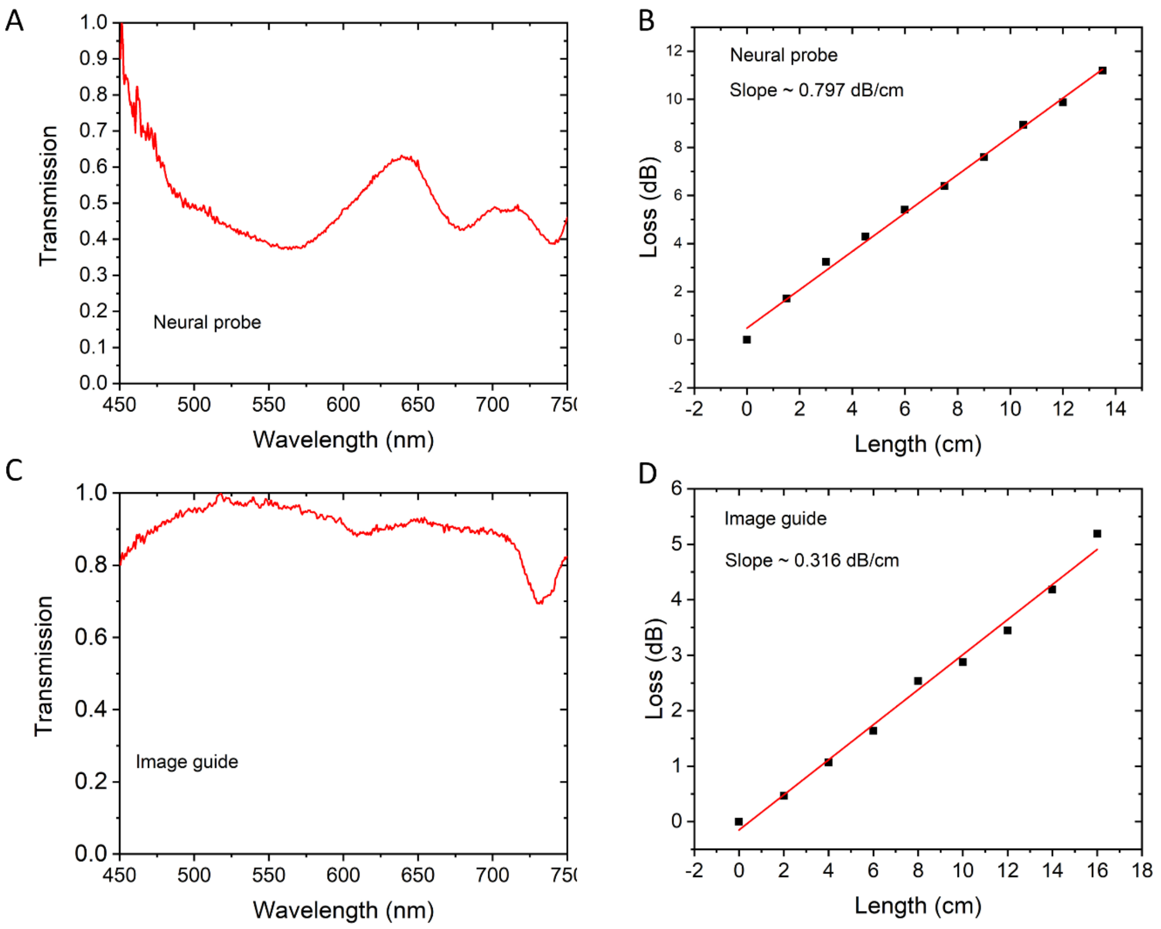


**Fig. S7 Characterization of optical properties of neural probe fiber (fiber F1) and image guide fiber.** (**A**) The optical transmission spectrum obtained at a wavelength range of 450 – 750 nm for neural probe fiber (normalized to the maximum value). (**B**) The transmission loss at a wavelength of 473 nm measured using cut-back method is 0.797 dB/cm ($R^2$=0.99592) for neural probe fiber. (**C**) The optical transmission spectrum obtained at a wavelength range of 450 – 750 nm for image guide fiber (normalized to the maximum value). (**D**) The transmission loss at a wavelength of 615 nm measured using cut back method is 0.316 dB/cm ($R^2$=0.9915) for image guide fiber.
